# Rafting a waterfall: Artificial selection for collective composition can succeed or fail depending on the initial and target values

**DOI:** 10.1101/2023.03.07.531234

**Authors:** Juhee Lee, Wenying Shou, Hye Jin Park

## Abstract

Collectives, such as microbial communities, can perform functions beyond the capability of individual members. Enhancing these collective functions through artificial selection, however, presents significant challenges. Here, we explore the ‘rafting-a-waterfall’ phenomenon, a metaphor illustrating how the success in achieving a target population composition in microbial collectives depends on both the target characteristics and initial conditions. Specifically, collectives comprising fast-growing (F) and slow-growing (S) individuals were grown for a period of “maturation” time, and the collective with S-frequency closest to the target value is chosen to “reproduce” (inoculate) offspring collectives. Such collective selection is dictated by two opposing forces: during collective maturation, intra-collective selection acts like a waterfall, relentlessly driving the S-frequency to lower values, while during collective reproduction, inter-collective selection resembles a rafter striving to reach the target frequency. Due to this model structure, maintaining a target frequency requires the continued action of inter-collective selection. Using simulations and analytical calculations, we show that intermediate target S frequencies are the most challenging, akin to a target within the vertical drop of a waterfall, rather than above or below it. This arises because intra-collective selection is the strongest at intermediate S-frequencies, which can overpower inter-collective selection. While achieving low target S frequencies is consistently feasible, attaining high target S-frequencies requires an initially high S-frequency — much like a raft that can descend but not ascend a waterfall. The range of attainable target frequencies depends on the initial population size of the collectives: as the population size in Newborn collectives increases, the region of achievable target frequency is reduced until no frequency is achievable. In contrast, the number of collectives under selection plays a less critical role. In scenarios involving more than two populations, the evolutionary trajectory must navigate entirely away from the metaphorical ‘waterfall drop.’ Our findings illustrate that the strength of intra-collective evolution is frequency-dependent, with implications in experimental planning.

## INTRODUCTION

Microbial collectives can carry out functions that arise from interactions among member species. These functions, such as waste degradation [1, 2], probiotics [3], and vitamin production [4], can be useful for human health and biotechnology. To improve collective functions, one can perform artificial selection (directed evolution) on collectives: Low-density “Newborn” collectives are allowed to “mature” during which cells proliferate and possibly mutate, and community function develops. “Adult” collectives with high functions are then chosen to reproduce, each seeding multiple offspring Newborns. Artificial selection of collectives have been attempted both in experiments [5–19] and in simulations [20–31], often with unimpressive outcomes.

One of the major challenges in selecting collectives is to ensure the inheritance of a collective function [32, 33]. Inheritance from a parent collective to offspring collectives can be compromised by changes in genotype and species compositions. During maturation of a collective, genotype compositions within each species can change due to intra-collective selection favoring fast-growing individuals, while species compositions can change due to ecological interactions. Furthermore, during reproduction of a collective, genotype and species compositions of offspring can vary stochastically from those of the parent.

Here, we consider the selection of collectives comprising two or three populations with different growth rates, and our goal is to achieve a target composition in the Adult collective. This is a common quest: whenever a collective function depends on both populations, collective function is maximised, by definition, at an intermediate frequency (e.g. too little of either population will hamper function [23]). Earlier work has demonstrated that nearly any target species composition can be achieved when selecting communities of two competing species with unequal growth rates [24, 34], so long as the shared resource is depleted during collective maturation [24]. In this case initially, both species evolved to grow faster, and the slower-growing species was preserved due to stochastic fluctuations in species composition during collective reproduction. Eventually, both species evolved to grow sufficiently fast to deplete the shared resource during collective maturation, and evolution in competition coefficients then acted to stabilize species ratio to the target value [24]. Regardless, earlier studies are often limited to numerical explorations, with prohibitive costs for a full characterization of the parameter space for such nested populations (population of collectives, and populations of mutants within a collective).

We mathematically examine the selection of composition in collectives consisting of populations growing at different rates. A selection cycle consists of three stages (Fig. 1). During collective maturation, intra-collective selection favors fast-growing individuals within a collective. At the end of maturation, inter-collective selection acts on collectives and favors those achieving the target composition. Finally during collective reproduction, offspring collectives sample stochastically from the parents, a process dominated by genetic drift. We made simplifying assumptions so that we can analytically examine the evolutionary tipping point between intra-collective and inter-collective selection. We show that this tipping point creates a “waterfall” effect which restricts not only which target compositions are achievable, but also the initial composition required to achieve the target. We also investigate how the range of achievable target composition is affected by the total population size in Newborns and the total number of collectives under selection. Finally, we show that the waterfall phenomenon extends to systems with more than 2 populations.

**Figure 1.**
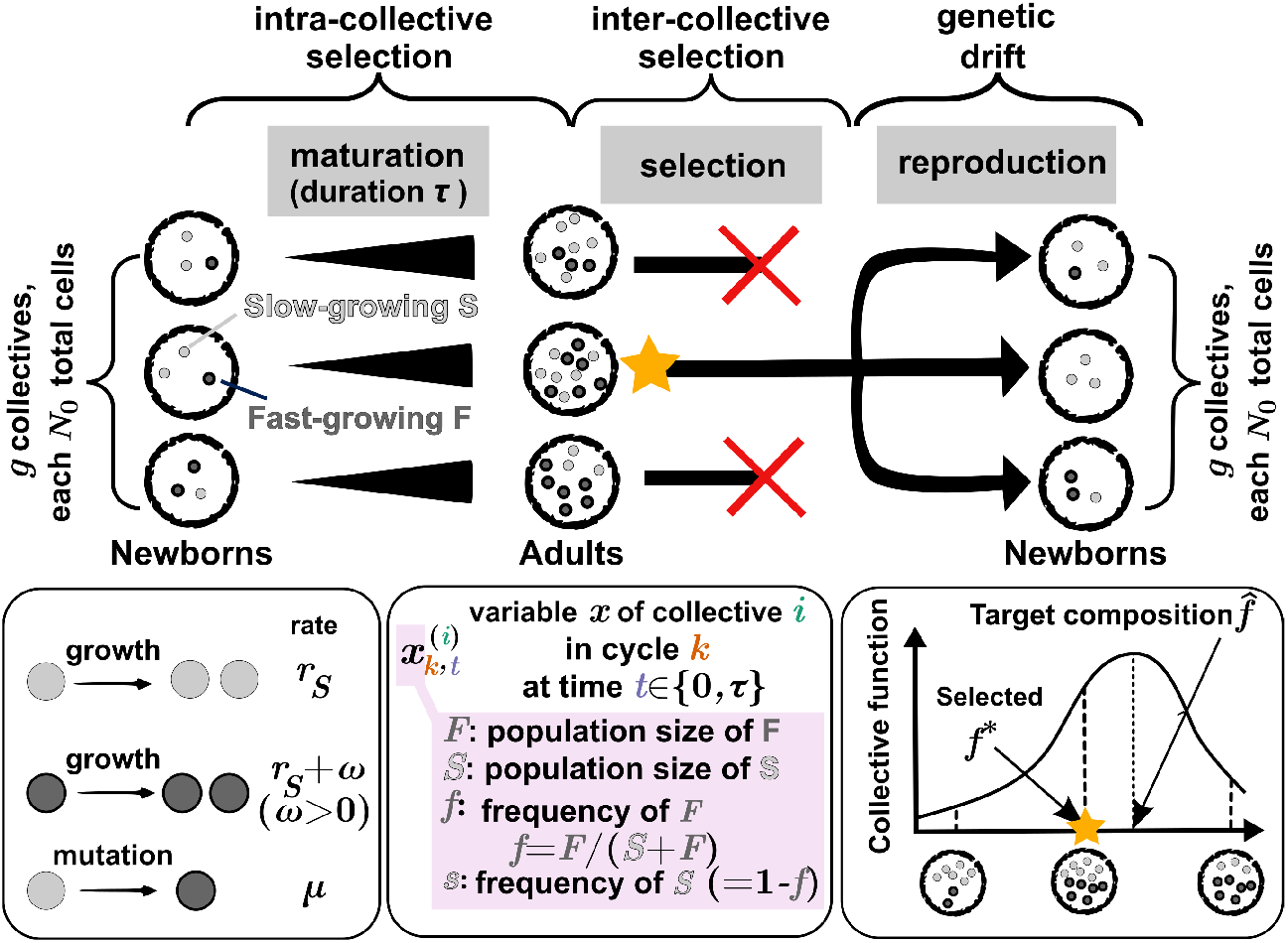
Schematic for artificial selection on collectives. Each selection cycle begins with a total of *g* Newborn collectives, each with *N*_0_ total cells of slow-growing S population (light gray dots) and fast-growing F population (dark gray dots). During maturation (over time *τ*), S and F cells divide at rates *r*_*S*_ and *r*_*S*_ + *ω* (*ω >* 0), respectively, and S mutates to F at rate *µ*. In the selection stage, the Adult collective with F frequency *f* closest to the target composition 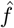 is chosen to reproduce *g* Newborns for the next cycle. Newborns are sampled from the chosen Adult (yellow star) with *N*_0_ cells per Newborn. The selection cycle is then repeated until the F frequency reaches a steady state, which may or may not be the target composition. To denote a variable *x* of *i*-th collective in cycle *k* at time *t* (0 ≤ *t* ≤ *τ*), we use notation 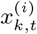 where *x* ∈ {*S, F, s, f*}. Note that time *t* = 0 is for Newborns and *t* = *τ* is for Adults.

## RESULTS AND DISCUSSIONS

To enable the derivation of an analytical expression, we have made the following simplifications. First, growth is always exponential, without complications such as resource limitation, ecological interactions between the two populations, or density-dependent growth. Thus, the exponential growth equation can be used. Second, we consider only two populations (genotypes or species): the fast-growing F population with size *F* and the slow-growing S population with size *S*. We do not consider a spectrum of mutants or species, since with more than two populations, an analytical solution becomes very difficult. Finally, the single top-functioning community is chosen to reproduce, which allows us to employ the simplest version of the extreme value theory (see section below for further justification.)

Our goal is to select for collective composition in terms of F frequency *f* = *F/*(*S* + *F*), or equivalently, S frequency *s* = 1 − *f*. More precisely, we want collectives such that after maturation time *τ, f* (*τ*) is as close to the target value 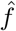 as possible (Fig. 1). Note that even if the target frequency has been achieved, since F frequency will always increase during maturation, inter-collective selection is required in each cycle to maintain the target frequency.

We will start with a complete model where S mutates to F at a nonzero mutation rate *µ*. We made this choice because it is more challenging to attain or maintain the target frequency when the abundance of fast-growing F is further increased via mutations. This scenario is encountered in biotechnology: an engineered pathway will slow down growth, and breaking the pathway (and thus faster growth) is much easier than the other way around. When the mutation rate is set to zero, the same model can be used to capture collectives of two species with different growth rates. We show that intermediate F frequencies or equivalently, intermediate S frequencies, are the hardest targets to achieve. We then show that similar conclusions hold when selecting for a target composition in collectives of more than two populations.

### Model structure

A selection cycle (Fig. 1) starts with a total of *g* Newborn collectives. At the beginning of cycle *k* (*t* = 0), each Newborn collective has a fixed total cell number 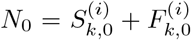 where 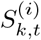 and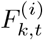, denote the numbers of S and F cells in collective *i* (1 ≤ *i* ≤ *g*) at time *t* (0 ≤ *t* ≤ *τ*) of cycle *k*. The average F frequency among the *g* Newborn collectives is *f*_*k*,0_, such that the initial F cell number in each Newborn is drawn from the binomial distribution Binom 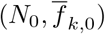.

Collectives are allowed to grow for time *τ* (‘Maturation’ in Fig. 1). During maturation, S and F grow at rates *r*_*S*_ and *r*_*S*_ + *ω* (*ω >* 0), respectively. If maturation time *τ* is too small, a matured collective (“Adult”) does not have enough cells to reproduce *g* Newborn collectives with *N*_0_ cells. On the other hand, if maturation time *τ* is too long, fast-growing F will take over. Hence we set the maturation time *τ* = ln(*g* +1)*/r*_*S*_, which guarantees sufficient cells to produce *g* Newborn collectives from a single Adult collective. At the end of a cycle, a single Adult with the highest function (with F frequency *f* closest to the target frequency 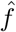) is chosen to reproduce *g* Newborn collectives each with *N*_0_ cells (‘Selection’ and ‘Reproduction’ in Fig. 1). Note that even though S and F do not compete for nutrients, they compete for space: because the total number of cells transferred to the next cycle is fixed, an overabundance of one population will reduce the likelihood of the other being propagated.

Collective function is dictated by the Adult’s F frequency *f*. Among all Adult collectives, the selected Adult is the one whose F frequency is closest to the target value, 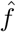.In contrast with findings from an earlier study [23], choosing top 1 is more effective than the less stringent “choosing top 5%”. In the earlier study, variation in the collective trait is partly due to nonheritable factors such as random fluctuations in Newborn biomass. In that context, a less stringent selection criterion proved more effective, as it helped retain collectives with favorable genotypes that might have exhibited suboptimal collective traits due to unfavorable nonheritable factors. However, since this study excludes nonheritable variations in collective traits, selecting the top 1 collective is more effective than selecting the top 5% (see Fig. 11 in Supplementary Information).

The selected Adult, with F frequency denoted as *f* ^*^, is then used to reproduce *g* offspring collectives, each with *N*_0_ total cells. The number of F cells in a Newborn follows a binomial distribution *B*(*N*_0_, *f* ^*^). By repeating the selection cycle, we aim to achieve and maintain the target composition 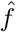.

**Table I.** Nomenclature.

| Variable | Representing |
| --- | --- |
| $S$ | Number of slower-growing (S) cells |
| $F$ | Number of faster-growing (F) cells |
| $s$ | Frequency of S cells, $s = S/(S + F)$ |
| $f$ | Frequency of F cells, $f = F/(S + F) = 1 - s$ |
| $r_S$ | Growth rate of S |
| $\omega > 0$ | Growth rate advantage of F over S |
| $\mu$ | Mutation rate from S to F |
| $g$ | Total number of collectives |
| $\tau$ | Maturation time |
| $N$ | Total cell numbers in a collective, $N = S + F$ |
| $N_0$ | Total number of cells in Newborn, or Newborn size |
| $\hat{s}, \hat{f}$ | Target frequency in $s$ or $f$ |
| $f^*$ | F frequency of the selected collective in a cycle |
| $f^L, f^H$ | Low and High thresholds of inaccessible $\hat{f}$ ($f^L < \text{inaccessible } \hat{f} < f^H$) |
| $R_\tau$ | Fold-growth of S cells over time $\tau$, $R_\tau = e^{r_S \tau}$ |
| $W_\tau$ | Fold ratio change of F cells over S cells over time $\tau$, $W_\tau = e^{\omega \tau}$ |

Overall our model considers mutational stochasticity, as well as demographic stochasticity in terms of stochastic birth and stochastic sampling of a parent collective by offspring collectives. Other types of stochasticity, such as environmental stochasticity and measurement noise, are not considered and require future research.

### The success of collective selection is constrained by the target composition, and sometimes also by the initial composition

Since intra-collective selection favors F, we expect that a higher target 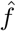 (a lower target *ŝ*) is easier to achieve in the sense that the absolute error *d* between the target frequency 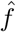 and the selected frequency averaged among independent simulations *(f* ^*^) is smaller than 0.05 (i.e.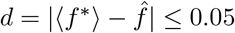).

We fixed *N*_0_, the total population size of a Newborn, to 1000, and obtained selection dynamics for various initial and target F frequencies by implementing stochastic simulations (Sec. I in Supplementary Information). If the target 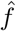 is high (e.g. 0.9, Fig. 2**a** magenta), selection is successful (computed absolute errors in Supplementary Information Fig. 4): regardless of the initial frequency, *f* ^*^ of the chosen collective eventually converges to the target 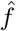 and stays around it. In contrast, without collective-level selection (e.g. choosing a random collective to reproduce), F frequency increases until F reaches fixation (Supplementary information Fig. 3**b**).

**Figure 2.**
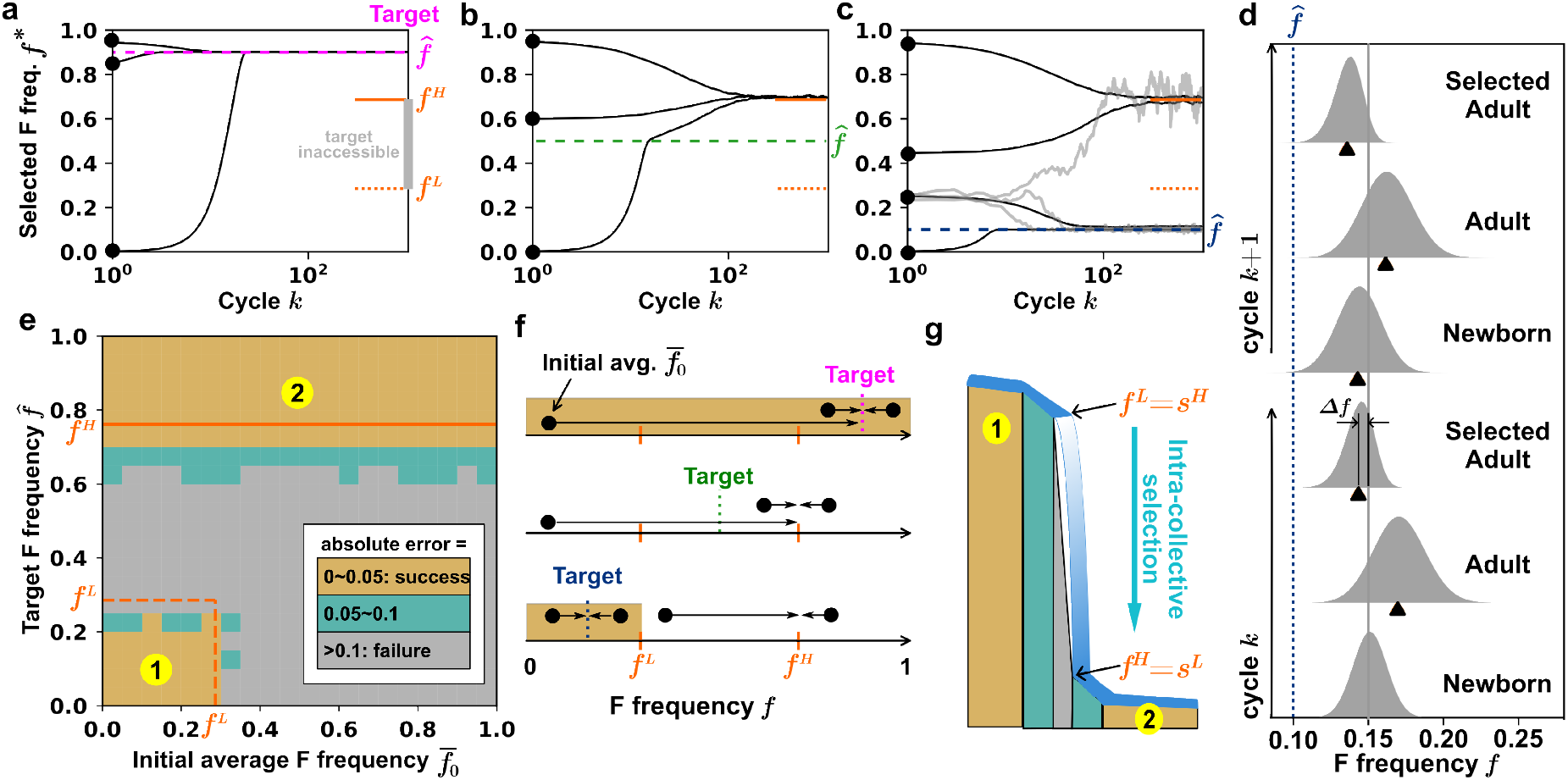
Initial and target compositions determine the success of artificial selection on collectives. **(a-c)** F frequency of the selected Adult collective (*f* ^*^) over cycles at different target 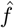 values (dashed lines). 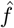 between *f* ^*L*^ and *f* ^*H*^ (orange dotted and solid line segments) is inaccessible where selection will fail. **a** A high target F frequency (e.g. 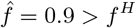; magenta dotted line) can be achieved from any initial frequency (black dots). **b** An intermediate target frequency (e.g. 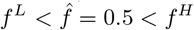; green dotted line) is never achievable, as all initial conditions converge to near *f* ^*H*^. **c** A low target frequency (e.g.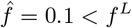) is achievable, but only from initial frequencies below *f* ^*L*^. For initial frequencies at *f* ^*L*^, stochastic outcomes (grey curves) are observed: while some replicates reached the target frequency, others reached *f* ^*H*^. For parameters, we used S growth rate *r*_*S*_ = 0.5, F growth advantage *ω* = 0.03, mutation rate *µ* = 0.0001, maturation time *τ*≈ 4.8, and *N*_0_ = 1000. The number of collectives *g* = 10. Each black line is averaged from independent 300 realizations. **d** Inter-collective selection opposes intra-collective selection. We plot probability density distributions of F frequency *f* during two consecutive cycles when selection is successful. Data correspond to cycles 31 and 32 from the second lowest initial point in **c.** Δ*f* is a selection progress (see Box.). Black triangle: median. **e** Two accessible regions (gold). Either high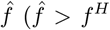 ; region 2) or low 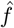 starting from low initial *f* (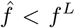and 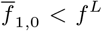 ; region 1) can be achieved. We theoretically predict (by numerically integrating Eq. (1)) *f* ^*H*^ (orange solid line) and *f* ^*L*^ (orange dotted line), which agree with simulation results (gold regions). **f** Example trajectories from initial compositions (black dots) to the target compositions (dashed lines). The gold areas indicate the region of initial frequencies where the target frequency can be achieved. **g** The tension between intra-collective selection and inter-collective selection creates a “waterfall” phenomenon. See the main text for details.

**Figure 3.**
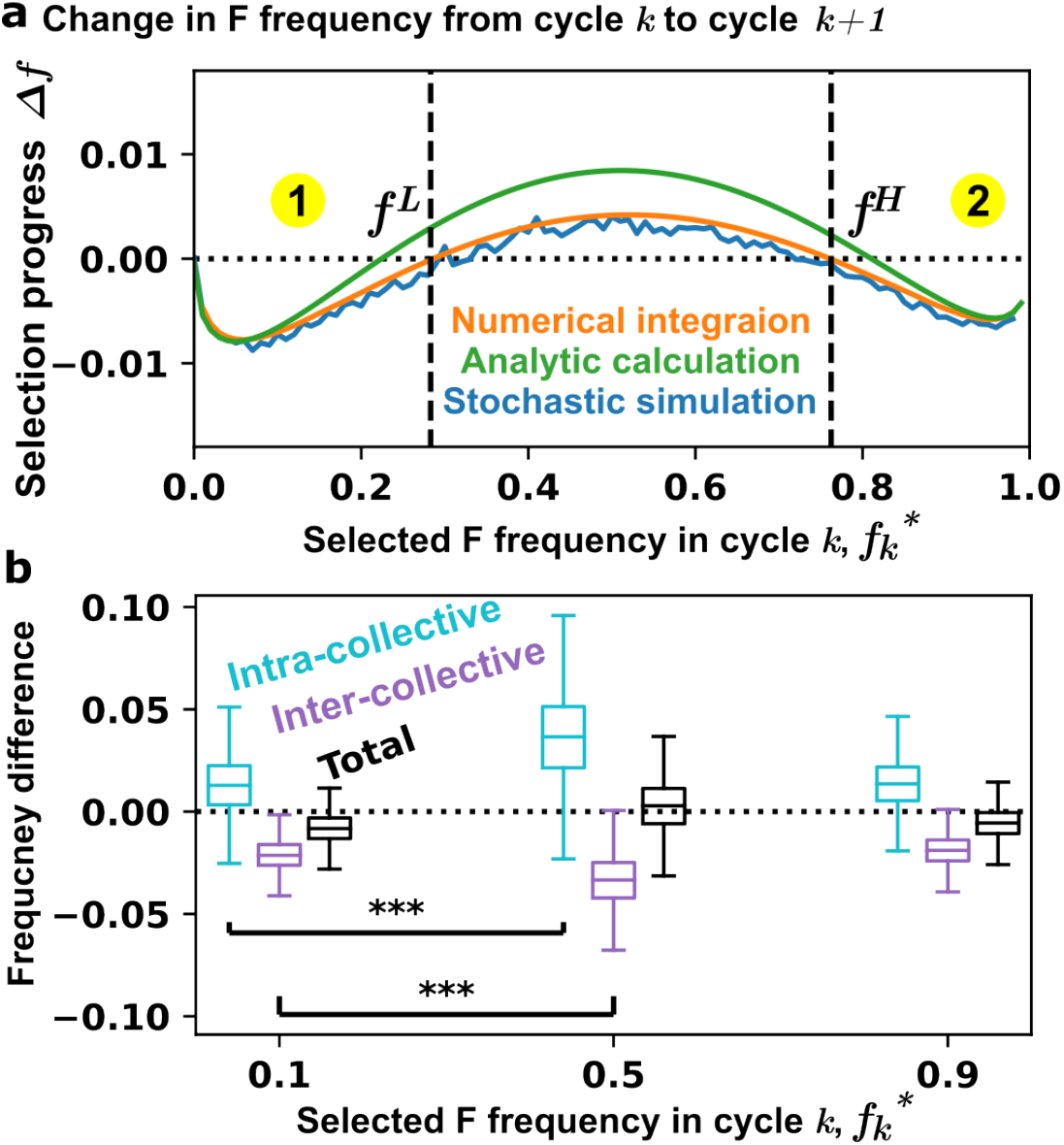
Intra-collective selection and inter-collective selection jointly set the boundaries for selection success. **a** The change in F frequency over one cycle. When 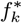 is sufficiently low or high, inter-collective selection can lower the F frequency to below 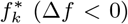.The points where Δ*f* = 0 (in the orange line) are denoted as *f* ^*L*^ and *f* ^*H*^, corresponding to the boundaries in Fig. 2. **b** The distributions of frequency differences obtained by 1000 numerical simulations. The cyan, purple, and black box plots respectively indicate the changes in F frequency after intra-collective selection (the mean frequency among the 100 Adults minus the mean frequency among the 100 Newborns during maturation), after inter-collective selection (the frequency of the 1 selected Adult minus the mean frequency among the 100 Adults), and over one selection cycle (the frequency of the selected Adult of one cycle minus that of the previous cycle). The box ranges from 25% to 75% of the distribution, and the median is indicated by a line across the box. The upper and lower whiskers indicate maximum and minimum values of the distribution. ***: *P <* 0.001 in an unpaired *t*-test.

**Figure 4.**
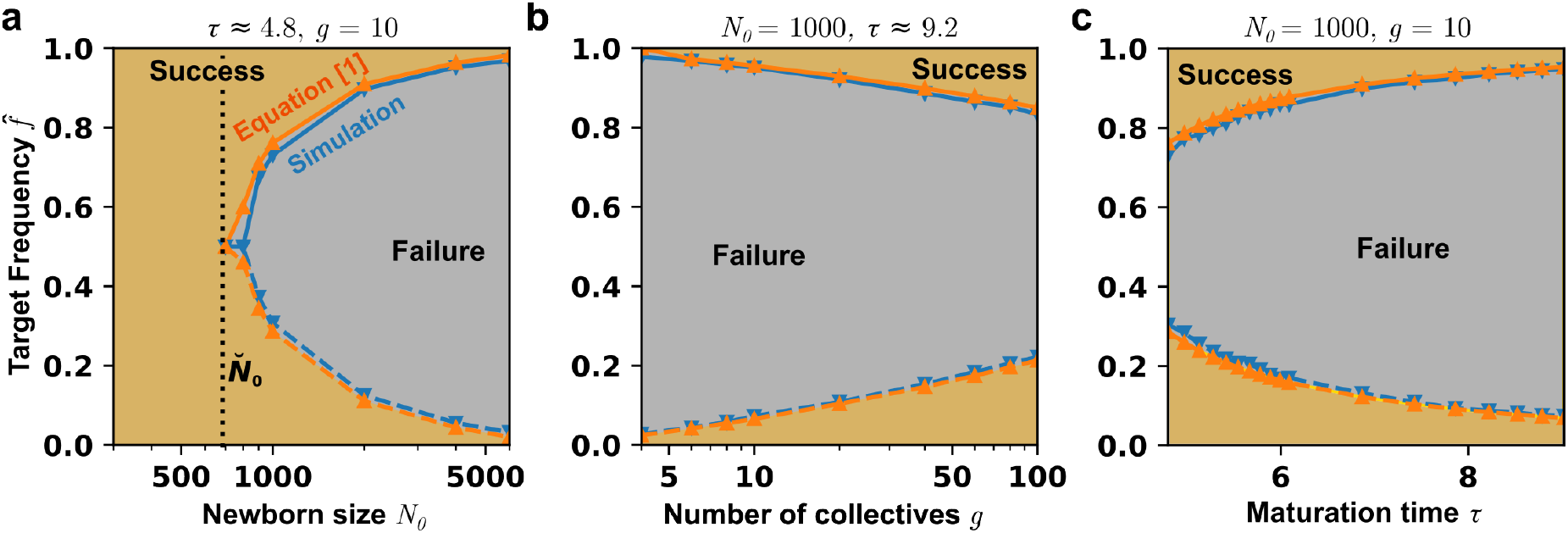
Expanding the success region for artificial collective selection. **a** Reducing the population size in Newborn *N*_0_ expands the region of success. In gold area, the probability that 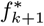 becomes smaller than 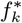 in a cycle is more than 50%. We used *g* = 10 and *τ* ≈ 4.8. Figures 2-3 correspond to *N*_0_ = 1000. Black dotted line indicates the critical Newborn size 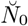,below which all target frequencies can be achieved. **b** Increasing the total number of collectives *g* also expands the region of success, although only slightly. We used a fixed Newborn size *N*_0_ = 1000. The maturation time *τ* (*τ* = log(100)*/r*_*S*_≈ 9.2) is set to be long enough so that an Adult can generate at least 100 Newborns. **c** Increasing the maturation time *τ* shrinks the region of success. We used a fixed Newborn size *N*_0_ = 1000 and number of collectives *g* = 10.

In contrast, an intermediate target frequency (e.g. 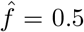 ; Fig. 2**b** green) is never achievable. High initial F frequencies (e.g. 0.6 and 0.95) decline toward the target, but stabilize at the “highthreshold” *f* ^*H*^ (∼ 0.7, solid orange line segment in Fig. 2**a-c**) above the target. Low initial F frequencies (e.g. 0) increase toward the target, but then overshoot and stabilize at the *f* ^*H*^ value.

If the target frequency is low (e.g.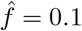; Fig. 2**c** dark blue), artificial selection succeeds when the initial frequency is below the “lower-threshold” *f* ^*L*^ (dotted orange line segment in Fig. 2**a-c**). Initial F frequencies above *f* ^*L*^ (e.g. 0.45 and 0.95) converge to *f* ^*H*^ instead. Initial F frequencies near *f* ^*L*^ display stochastic trajectories, converging to either *f* ^*H*^ or the target.

To achieve target 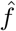,inter-collective selection must overcome intra-collective selection. We can visualize the distributions of *f* over two consecutive cycles (bottom to top, Fig. 2**d**) where *f* started above target 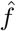.When Newborns matured into Adults, the distribution of *f* up-shifted due to intra-collective selection. The distribution of *f* was then down-shifted toward the target due to inter-collective selection. If the magnitude of down-shift exceeded that of up-shift, progress toward the target was made. During reproduction, the distribution of *f* retained the same mean but became broader due to stochastic sampling by the Newborns from their parent.

In summary, two regions of target frequencies are “accessible” in the sense that a target value in the region can be reached (gold in Fig. 2**e, f**): (1) target frequencies above 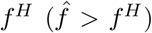 or (2) target frequencies below 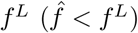 and starting at an average frequency below 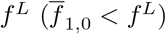.

### Intra-collective evolution is the fastest at intermediate F frequencies, creating the “waterfall” phenomenon

To understand what gives rise to the two accessible regions, we calculated Δ*f*, the selection progress in F frequency over two consecutive cycles (Box 1, Eq. (2)). The solution (Fig. 3**a**, green) has the same shape as results from numerically integrating Eq. (1) (Fig. 3**a**, orange) and from stochastic simulations (Fig. 3**a**, blue).

If Δ*f* is negative, then inter-collective selection will succeed in countering intra-collective selection and reducing *f* toward the target. Δ*f* is negative if the selected 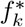 is low or high, but not if it is intermediate between *f* ^*L*^ and *f* ^*H*^ (Fig. 3**a**). This is because *f* increase during maturation is the most drastic when Newborn *f* is intermediate (Fig. 3**b**), for intuitive reasons: when Newborn *f* is low, the increase in *f* will be minor; when Newborn *f* is high, the fitness advantage of F over the population average is small and hence the increase is also minor. Thus, when Newborn F frequency is intermediate, inter-collective selection may not be able to counter intra-collective selection (Fig. 3**b** and Fig. 6**a** in Supplementary Information). Not surprisingly, similar conclusions are derived where S and F are slow-growing and fast-growing species which cannot be converted through mutations (Sec. IV and Fig. 8 in Supplementary Information).

Thus, inter-collective selection is akin to a raftman rowing the raft to a target, while intra-collective selection is akin to a waterfall. This metaphor is best understood in terms of S frequency *s* = 1 − *f*. The lower-threshold *f* ^*L*^ corresponds to higher-threshold in *s*^*H*^ = 1 − *f* ^*L*^, while higher-threshold *f* ^*H*^ corresponds to lower-threshold in *s*^*L*^ = 1 − *f* ^*H*^. Intra-collective selection is akin to a waterfall, driving the S frequency *s* from high to low (Fig. 2**g**). Intra-collective selection acts the strongest when *s* is intermediate (*s*^*L*^ *< s <* s^*H*^), similar to the vertical drop of the fall. Intra-collective selection acts weakly at high (*> s*^*H*^) or low (*< s*^*L*^) *s*, similar to the gentle sloped upper and lower pools of the fall (regions 1 and 2 of Fig. 2**e, g**). Thus, an intermediate target frequency can be impossible to achieve − a raft starting from the upper pool will be flushed down to *s*^*L*^ (*f* ^*H*^), while a raft starting from the lower pool cannot go beyond *s*^*L*^ (*f* ^*H*^). In contrast, a low target S frequency (in the lower pool) is always achievable. Finally, a high target S frequency (in the upper pool) can only be achieved if starting from the upper pool (as the raft can not jump to the upper pool if starting from below).

#### Box 1

Changes in the distribution of F frequency *f* after one cycle

We consider the case where 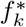,the F frequency of the selected Adult at cycle *k*, is above the target value 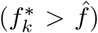.This case is particularly challenging because intra-collective evolution favors fast-growing F and thus will further increase *f* away from the target. From 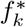,Newborns of cycle *k* + 1 will have *f* fluctuating around 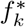,and after they mature, the minimum *f* is selected 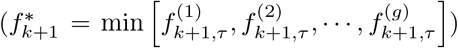. If the selected composition at cycle *k* + 1 can be reduced compared to that of cycle *k* (i.e.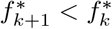), the system can evolve to the lower target value.

To find 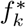 values such that 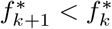,we used the median value of the conditional probability distribution Ψ of 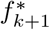 (the F frequency of the selected Adult at cycle *k* + 1 being *f*) given the selected 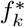 at cycle *k* (mathematical details in Sec. II in Supplementary Information). If the median value (Median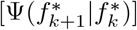) is smaller than 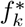,then selection will likely be successful since the selected Adult in cycle *k* + 1 has more than 50% chance to have a reduced F frequency compared to cycle *k*.

There are two points where the median values are the same as 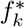 (Fig. 3**a**), which are assigned as lower-threshold (*f* ^*L*^) and higher-threshold (*f* ^*H*^).

Following the extreme value theory, the conditional probability density function 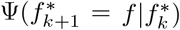 of 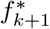 (the F frequency of the selected Adult at cycle *k* + 1 being *f*) given the selected 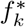 at cycle *k* is

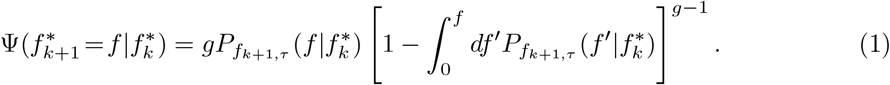

Equation (1) can be described as the product between two terms related to probability: (i) 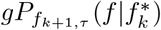 describes the probability density that any one of the *g* Adult collectives achieves *f* given 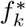,and (ii) 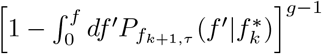 describes the probability that all other *g* − 1 collectives achieve frequencies above *f* and thus not selected.

Since computing the exact formula of Adults’ *f* distribution in cycle *k* + 1 is hard, we approximate it as Gaussian with mean 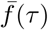 and variance 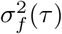.The Gaussian approximation on Eq. (1) requires sharp Gaussian distributions of *S*(*τ*) and *F* (*τ*) (i.e. 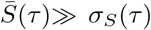 and 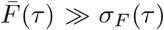).Compared to Gaussian, the exact *S*(*τ*) (negative binomial) distribution and *F* (*τ*) (Luria-Delbrück) distribution are right-skewed and heavy-tailed. However these problems are alleviated when the initial numbers of *S* and *F* cells are not small (on the order of 100). Additionally, the sharpness of distributions could be achieved (see Fig. 1 in Supplementary Information).

To obtain an analytical solution of the change in *f* over one cycle, we first assume that in a Newborn collective, the number of S cells is distributed as Gaussian with mean 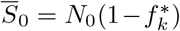 and variance 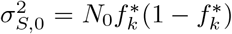.Then, the number of F cells, *F*_0_ = *N*_0_ − *S*_0_, is distributed as Gaussian with mean 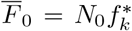 and variance 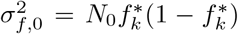.From these, we can calculate for Adult collectives the mean and variance of population sizes *F* (*τ*) (i.e. 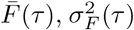) and *S*(*τ*) (i.e.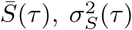) (mathematical details in Sec. I in Supplementary Information). This task is simplified by the exponential growth of S and 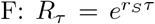 describes the foldgrowth of S over maturation time *τ*, and since *ω* is the fitness advantage of F over S, *W*_*τ*_ = *e*^*ωτ*^ describes the fold change of F/S over time *τ*. From 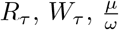 (mutation rate scaled with the fitness difference), 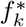 (F frequency in the selected collective at cycle *k*), *N*_0_ (Newborn size), 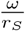 (relative fitness advantage), we can calculate the mean and variance of F frequency among the Adults of *k* + 1 cycle 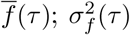,detailed formula in Eqs. (40) and (41) in Supplementary Information).

Selection progress - the difference between the median value of the conditional probability distribution 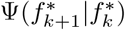 and the selected frequency of 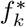 (Sec. II in Supplementary Information) - can be expressed as:

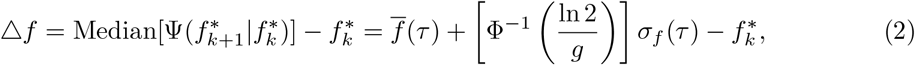

where Φ^−1^(…) is the inverse cumulative function of standard normal distribution (see main text for an example). We chose median because compared to mean, it is easier to get analytical expression since Φ^−1^(…) is known in a closed form. Regardless, using median generated results similar to simulations (Fig. 7 in Supplementary Information). As expected, selection progress Δ*f* is governed by both the mean 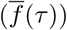 and the variation (*σ*_*f*_ (*τ*)) in *f* among Adults.

When 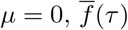 and *σ*_*f*_ (*τ*) can be simplified to:

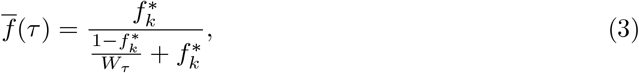

and

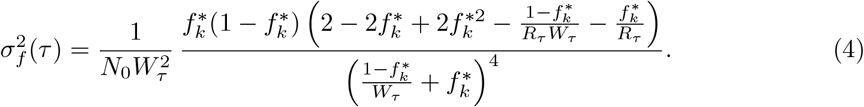

In the limit of small 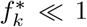,Equation (3) becomes 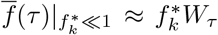 while Equation (4) becomes 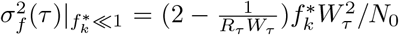.Thus, both Newborn size (*N*_0_) and fold-change in F/S during maturation (*W*_*τ*_) are important determinants of selection progress.

### Manipulating experimental setups to expand the achievable target region

In Eq. (2) (Box 1), selection progress Δ*f* depends on the total number of collectives under selection (*g*). Δ*f* also depends on the mean and the standard deviation of Adult F frequency 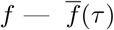 and *σ*_*f*_ (*τ*). Eqs. (3) and (4) (Box 1) provide simplified expressions of 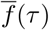 and *σ*_*f*_ (*τ*) when mutation rate *µ* has been set to 0. When mutation rate *µ* is not zero (Eqs. (40) and (41) in Section II of Supplementary Information), selection progress is additionally influenced by 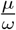 (mutation rate *µ* scaled with fitness difference *ω*).

Our goal is to make Δ*f* as negative as possible so that any increase in *f* during collective maturation may be reduced. From Eq. (2) (Box 1), a small *f* (*τ*) will facilitate collective-level selection. Additionally, a large *σ*_*f*_ (*τ*) will also facilitate collective-level selection due to negative 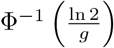.Note that since 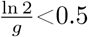 for 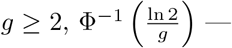 corresponding to the number *y* such that the probability of a standard normal random variable being less than or equal to *y* is 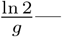 is negative.

From Eq. (4) (Box 1), *σ*_*f*_ (*τ*) will be large if Newborn size *N*_0_ is small. Indeed, as Newborn size *N*_0_ declines, the region of achievable target frequency expands (larger gold area in Fig. 4**a**). If the Newborn size *N*_0_ is sufficiently small (e.g. ≤ 700 in our parameter regime), any target frequency can be reached. An analytical approximation of the maximal Newborn size permissible for all target frequencies is given in section III in Supplementary Information.

From Eqs. (3) and (4) (Box 1), maturation time *τ* affects 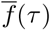 through 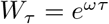 (the fold change in F/S over *τ*), and affects 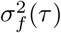 additionally through 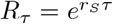 (fold-growth of S over *τ*). Longer *τ* increases 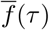 and is thus detrimental to selection progress. The relationship between *σ*_*f*_ (*τ*) and *τ* is not monotonic (Fig. 6**c** in Supplementary Information), meaning that an intermediate value of *τ* is the best for achieving large *σ*_*f*_ (*τ*). However, the effect of 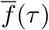 dominates that of *σ*_*f*_ (*τ*) and therefore, the region of success monotonically reduces with longer maturation time (Fig. 4**c**). Similarly, 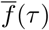 will be small if *ω* (fitness advantage of F over S) is small. Indeed, as *ω* become larger, the region of success becomes smaller (Fig. 9 in Supplementary Information).

*g*, the number of collectives under selection, also affects selection outcomes. As *g* increases, the value of 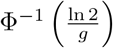 becomes more negative, and so does Δ*f* - meaning collective-level selection will be more effective. Intuitively, with more collectives, the chance of finding a *f* closer to the target is more likely. Thus, a larger number of collectives broadens the region of success (Fig. 4**b**). However, the effect of *g* is not dramatic. To see why, we note that the only place that *g* appears is Eq. (2) in 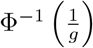. When *g* becomes large, 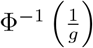 is asymptotically expressed as 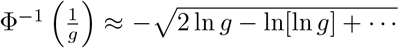 ln (Section II in Supplementary Information)[35], and thus does not change dramatically as *g* varies.

### The waterfall phenomenon in a higher dimension

To examine the waterfall effect in a higher dimension, we investigate a three-population system where a faster-growing population (FF) grows faster than the fast-growing population (F) which grows faster than the slow-growing population (S) (Fig. 5**a** and Sec. VIII in Supplementary Information). In the three-population case, the evolutionary trajectory travels in a two-dimensional plane. A target population composition can be achieved if inter-collective selection can sufficiently reduce the frequencies of F as well as FF (accessible region, gold in Fig. 5**b**).

**Figure 5.**
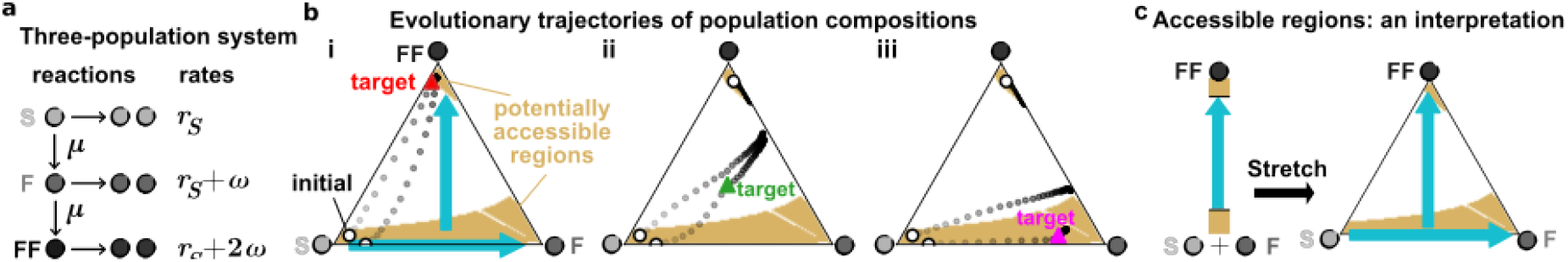
In higher dimensions, the success of artificial selection requires the entire evolutionary trajectory remaining in the accessible region. **a** During collective maturation, a slow-growing population (S) (with growth rate *r*_*S*_ ; light gray) can mutate to a fast-growing population (F) (with growth rate *r*_*S*_ + *ω*; medium gray), which can mutate further into a faster-growing population (FF) (with growth rate *r*_*S*_ + 2*ω*; dark gray). Here, the rates of both mutational steps are *µ*, and *ω >* 0. **b** Evolutionary trajectories from various initial compositions (open circles) to various targets. Intra-collective evolution favors FF over F (vertical blue arrow) over S (horizontal blue arrow). The accessible regions are marked gold (see Sec. I in Supplementary Information). We obtain final compositions starting from several initial compositions while aiming for different target compositions in i, ii, and iii. The evolutionary trajectories are shown in dots with color gradients from the initial to final time. (i) A target composition with a high FF frequency is always achievable. (ii) A target composition with intermediate FF frequency is never achievable. (iii) A target composition with low FF frequency is achievable only if starting from an appropriate initial composition such that the entire trajectory never meanders away from the accessible region. The figures are drawn using mpltern package [36]. **c** The accessible region in the three-population problem is interpreted as an extension of the two-population problem. First the accessible region between FF and S+F is given, and then the S+F region is stretched into S and F.

From numerical simulations, we identified two accessible regions: a small region near FF and a band region spanning from S to F (gold in Fig. 5**b** i). Intuitively, the rate at which FF grows faster than S+F is greater than the rate at which F grows faster than S (see section VIII in Supplementary Information). Thus, the problem can initially be reduced to a two-population problem (i.e. FF versus F+S; Fig. 5**c** left), and then expanded to a three-population problem (Fig. 5**c** right).

Similar to the two-population case, targets in the inaccessible region are never achievable (Fig. 5**b** ii), while those in the FF region are always achievable (Fig. 5**b** i). Strikingly, a target composition in an accessible region may not be achievable even when the initial composition is within the same region: once the composition escapes the accessible region, the trajectory cannot return back to the accessible region (Fig. 5**b**iii, the leftmost initial condition). However, if the initial position is closer to the target in the accessible region, the target becomes achievable (Fig. 5**b** iii, initial condition near the bottom). Note that here, selection outcome is path-dependent in the sense of being sensitive to initial conditions. This phenomenon is distinct from hysteresis where path-dependence results from whether a tuning parameter is increased or decreased.

In conclusion, we have investigated the evolutionary trajectories of population compositions in collectives under selection, which are governed by intra-collective selection (which favors fast-growing populations) and inter-collective selection (which, in our case, strives to counter fast-growing populations). Intra-collective selection has the strongest effect at intermediate frequencies of faster-growing populations, potentially creating an inaccessible region analogous to the vertical drop of a waterfall. High and low target frequencies are both accessible, analogous to the lower- and the upper-pools of a waterfall, respectively. A less challenging target (high 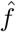; low *ŝ*) is achievable from any initial position. In contrast, a more challenging target (low 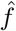; high *ŝ*) is only achievable if the entire trajectory is contained within the region, similar to a raft striving to reach a point in the upper-pool must start at and remain in the upper pool. Our work suggests that the strength of intra-collective selection is not constant, and that strategically choosing an appropriate starting point can be essential for successful collective selection.

## METHODS

### Stochastic simulations

A selection cycle is composed of three steps: maturation, selection, and reproduction. At the beginning of the cycle *k*, a collective *i* has 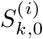 slow-growing cells and 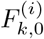 fast-growing cells. At the first cycle, mean F frequency of collectives are set to be 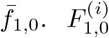 is sampled from the binomial distribution with mean 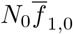. Then, 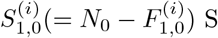 cells are in the collective *i*. In the maturation step, we calculate 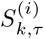 and 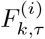 by using stochastic simulation. We can simulate the division and mutation of each individual cell stochastically by using tau-leaping algorithm [37, 38](see Fig. 3**a** and **b** in Supplementary Information). However, individual-based simulations require a long computing time. Instead, we randomly sample 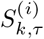 and 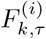 from the joint probability density distribution 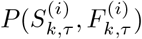.To obtain 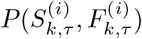,we solve the master equation which describes a time evolution of probability distribution 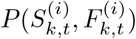 under the random processes (see Sec. I in Supplementary Information). We assumed that 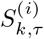 and 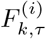 are independent (as S and F populations grow independently without ecological interactions), and thus 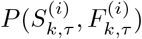 is product of two probability density functions 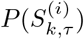 and 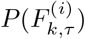.Each distribution follows a Gaussian distribution,with the mean and variance numerically obtained from ordinary differential equations derived from the master equation (see Sec. I in Supplementary Information). We choose the collective with the closest frequency to the target 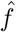 to generates *g* Newborns. The number of F cells is sampled from the binomial distribution with the mean of 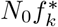.We start a new cycle with those Newborn collectives. Then, the number of S cells in a collective *i* is 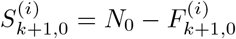.

### Analytical approach to the conditional probability

The conditional probability distribution 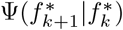 of observing 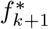 at a given 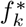 is calculated by the following procedure. Given the selected collective in cycle *k* with 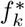,the collective-level reproduction proceeds by sampling *g* Newborn collectives with *N*_0_ cells in cycle *k* + 1. Each Newborn collective contains certain F numbers 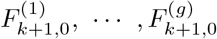 at the beginning of the cycle *k* + 1, which can be mapped into 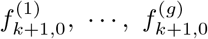 with the constraint of *N*_0_ cells. If the number of cells in the selected collective is large enough, the joint conditional distribution function 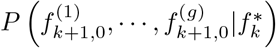 is well described by the product of *g* independent and identical Gaussian distribution 𝒩 (*µ, σ*^2^). So we consider the frequencies of *g* Newborn collectives as *g* identical copies of the Gaussian random variable *f*_*k*+1,0_. The mean and variance of *f*_*k*+1,0_ are given by 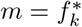 and 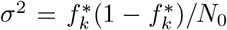.Then, the conditional probability distribution function of *f*_*k*+1,0_ being *ζ* is given by

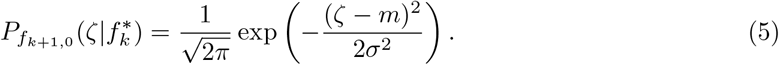

After the reproduction step, the Newborn collectives grow for time *τ*. The frequency is changed from the given frequency *ζ* to *f* by division and mutation processes. We assume that the frequency *f* of an Adult is also approximated by a Gaussian random variable 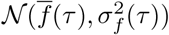.The mean 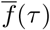 and variance 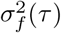 are calculated by using means and variances of *S* and *F* (see Sec. II in Supplementary Information.) Since 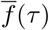 and 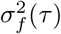 also depend on *ζ*, the conditional probability distribution function of *f*_*k*+1,*τ*_ being *f* is given by

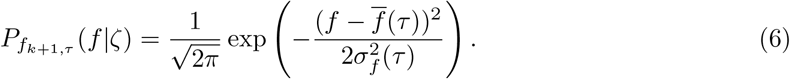

The conditional probability distribution of an Adult collective in cycle *k* + 1 (*f*_*k*+1,*τ*_) to have frequency *f* at a given 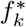 is calculated by multiplying two Gaussian distribution functions and integrating overall *ζ* values, which is given by

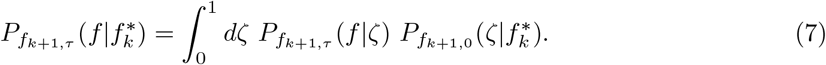

Since we select the minimum frequency 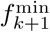 among *g* identical copies of *f*_*k*+1,*τ*_, the conditional probability distribution function of 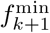 follows a minimum value distribution, which is given in Eq. (1). Here, for the case of 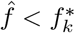,the selected frequency 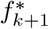 is the minimum frequency 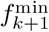. So we have 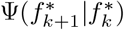 by replacing 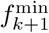 with 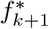.

We assume that the conditional probability distribution in Eq. (7) follows a normal distribution, whose mean and variances are describe by Eq. (40) and Eq. (41). Then, the extreme value theory [39] estimates the median of the selected Adult by

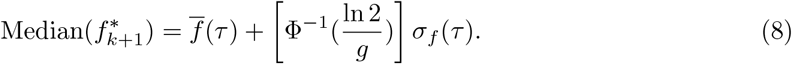

The selection progress Δ*f* in Eq. (2) is obtained by subtracting 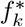 from Eq. (8).

## DATA AND SOURCE CODE AVAILABILITY

Data and source code of stochastic simulations are available in https://github.com/schwarzg/artificial_selection_collective_composition

## ACKNOWLEDGMENTS

J. Lee and H.J. Park were supported by the National Research Foundation of Korea grant funded by the Korea government (MSIT), Grant No. RS-2023-00214071 and RS-2024-00460958 and by an appointment to the JRG Program at the APCTP through the Science and Technology Promotion Fund and Lottery Fund of the Korean Government. W. Shou was supported by an Academy of Medical Sciences Professorship and a Royal Society Wolfson Fellowship. This was also supported by the Korean Local Governments–Gyeongsangbuk-do Province and Pohang City and INHA UNIVERSITY Research Grant. We thank Su-Chan Park, Li Xie, Alex Yuan, and Botond Major for constructive comments and discussions.

## Supplementary Information

In this supplementary information (SI), we present the mathematical details of all calculations in the main text, as well as additional figures. The SI is organized as follows: In section I, we provide the mathematical details of stochastic simulation of the selection cycle. In section II, we show the mathematical process to obtain the analytical expression of the selection progress in Eq.(2) in the main text. In section III, we provide the analytical approximation of the critical Newborn size. In section IV, we discuss a limiting case when mutation rate is zero. Section V provides the boundary of the region of success with different selective advantages. In section VI, we discuss specific scenarios under which mutation converts F to S. In section VII, we present an example in which more than one collective is selected for reproduction. In section VIII, we provide the mathematical details of the selection cycle for a three-population system. Finally, in section IX, we provide detailed derivations of equations from section I to section II.

### I STOCHASTIC SIMULATION OF THE SELECTION CYCLE

In the main text, we design a simple model of artificial selection on collectives. The selection cycle starts with *g* “Newborn” collectives which consist of two populations - slow-growing population (S) and fast-growing population (F). S mutate to F at a rate *µ*. The Newborns mature for a fixed time *τ*. The matured collective (“Adult”) with the highest function (with F frequency *f* closest to the target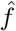) is chosen to reproduce *g* Newborn collectives each with *N*_0_ cells.

In our selection cycle, variation among collectives mainly resulted from demographic noises during cell birth, cell mutation, and collective reproduction. In this section, we provide details of the simulation.

#### Maturation

Here, we calculate the cell numbers during maturation. Each collective *i* (*i* = 1, …, *g*) has 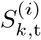 S cells and 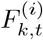 F cells where *k* is the cycle number and *t* indicates time (0 ≤ *t* ≤ *τ*). At the beginning of cycle *k* (*t* = 0), each Newborn collective has a total of 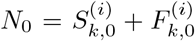 cells. The collectives are allowed to “mature” for *t* = *τ* during which S and F grow at rates *r*_*S*_ and *r*_*S*_ + *ω* (*ω >* 0), respectively. In this subsection, we ignore the cycle number index *k* and the collective index (*i*) for convenience. That is, we denote 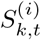 and 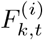 as *S*(*t*) and *F* (*t*), respectively.

We describe cell divisions of S and F cells and mutation from S to F with following chemical reaction rules:

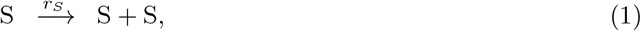

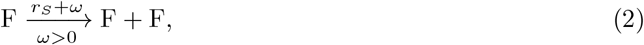

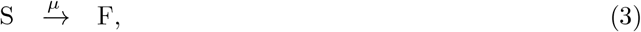

One can run an individual-based simulation by counting the number of events occurring during collective maturation via the tau-leaping algorithm [1, 2] to generate a sample trajectory of *S*(*t*) and *F* (*t*) for each collective. However, the individual-based simulation requires long computing times due to a large number of random events to be counted. Hence we used a ‘sampling method’ by sampling the numbers of S and F cells in collectives from a joint probability density distribution (jpdf) *P* (*S, F, t*) which denotes the probability density to have *S* number of S cells and *F* number of F cells at time *t* in the cycle. To do so, we require an analytical expression of *P* (*S, F, t*).

First, we assume that the chemical reactions in Eqs. (1)-(3) occur independently, and never occur simultaneously within a short time interval [*t, t* + *dt*). Then, the differential of *P* (*S, F, t*) with respect to time is given by

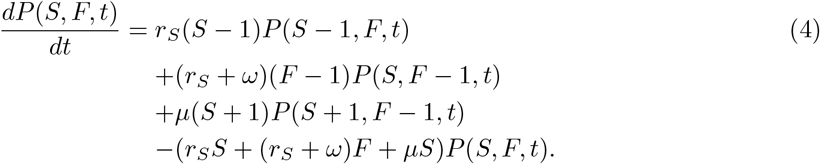

This master equation describes a probability density ‘flux’ at the state (*S, F*). The first term describes the scenario where a single birth event of a S cell happens during time interval [*t, t* + *dt*), which changes the collective’s composition from (*S*− 1, *F*) to (*S, F*). Similarly, the second term comes from a birth event of a F cell. The third term indicates the mutation event from (*S* +1, *F*− 1) to (*S, F*). The last term corresponds to the outflow of probability density by birth and mutation processes, which describes the changes from (*S, F*) to any other states.

Calculating the exact form of *P* (*S, F, t*) is not simple. Instead, we assume that the mutation rate is much smaller than the growth rates, and hence the correlation between *S* and *F* is sufficiently small. Additionally, S and F do not interact ecologically. Then, we can express *P* (*S, F, t*) as a product of two probability density functions (pdf) of *S*(*t*) and *F* (*t*), *P* (*S, F, t*) = *P* (*S, t*)*P* (*F, t*). We assume that each pdf of *S* and *F* can be approximated as Gaussian (𝒩), which is supported by the Central Limit Theorem and Fig. 1. In more detail, the cell numbers *S* and *F* are mainly determined by growth (Eqs. (1) and (2)), and also mutations (Eq. (3)). Even though the number of events would be different among different realizations, the mean numbers of events will follow Gaussian distributions. So, we can simply assume that the distributions of cell numbers also follow Gaussian distributions. This assumption requires that the distributions have insignificant skewness and no heavy tails, which we will numerically check afterwards. The pdfs of *S*(*t*) and *F* (*t*) are given by

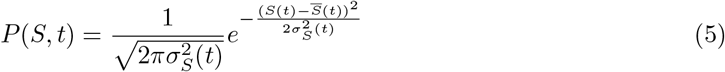

and

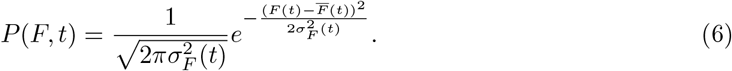

That is, *P* (*S, F, t*) is written as

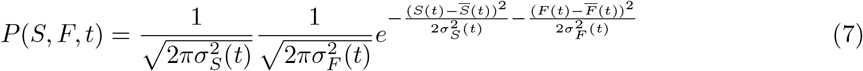

Now we need means (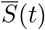 and 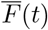) and variances (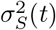 and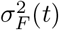) of S and F cell numbers to express the distribution analytically.

The means are defined by 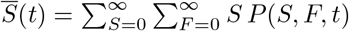 and 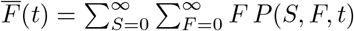.The differential equations for means are obtained by applying the definition to the master equation in Eq. (4), as

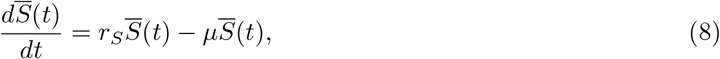

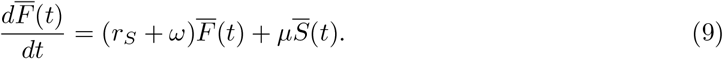

We assume that the mutation rate *µ* is much smaller than *r*_*S*_ and *ω*. By solving Eq. (8) and Eq. (9), the means 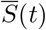 and 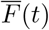 are given by

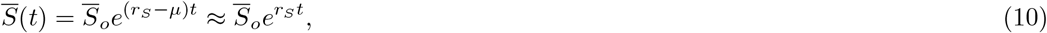

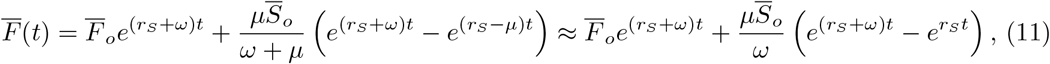

where 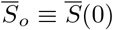 and 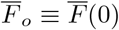 are the mean numbers of S and F cells at the beginning of cycle, *t* = 0. Note that the second term of Eq. (11) is consistent with previous studies [3]. Now we introduce factors 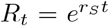 and *W*_*t*_ = *e*^*ωt*^ in Eqs. (10) and (11) in order to simplify the formula. *R*_*t*_ is the multiplying factor by which the S cell number increases after time *t. W*_*t*_ is the fold change in *F/S*. Then, we can rewrite

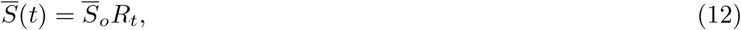

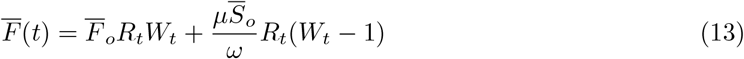

We define the second momenta of *S* and *F* as

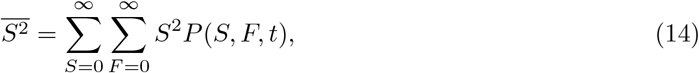

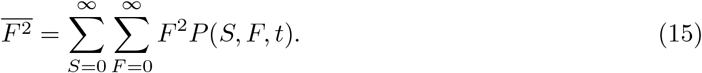

Then, the corresponding differential equations are given by

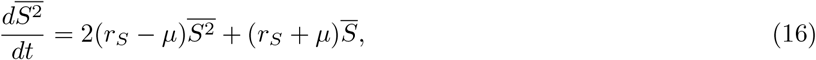

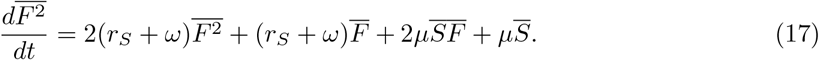

The solution of Eq. (16) is

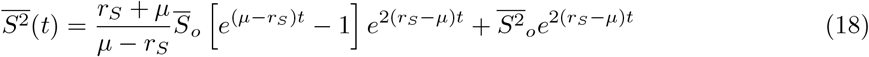

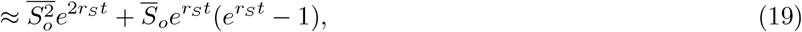

where 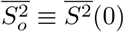 is the second moment of initial values. Thus, the variance 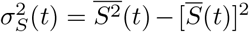 is

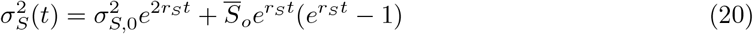

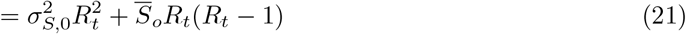

where 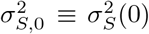 is a variance of S cell numbers at *t* = 0. In Eq. (17), we require 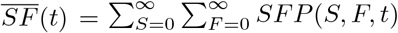 to calculate 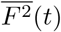.Eq. (4) provides a differential equation for 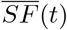 as

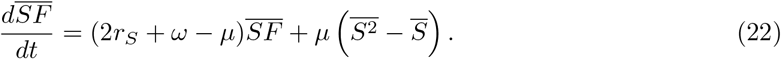

The solution of Eq. (22) is given by

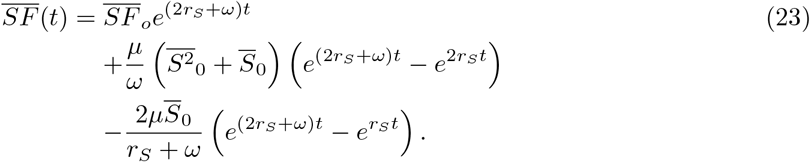

By using Eq. (23), the solution of Eq. (17) is given by

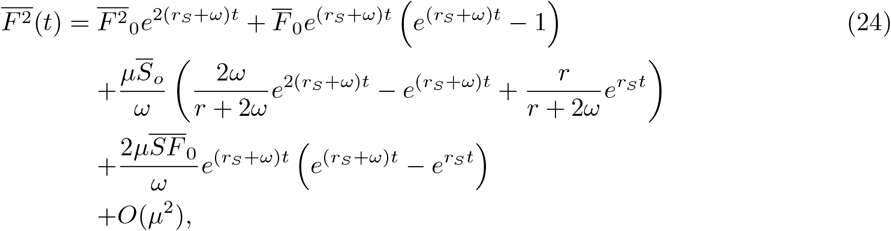

where 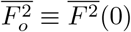 is the second moment of initial values. Thus, the variance 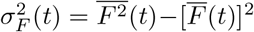 is given, up to the order of *µ*, by

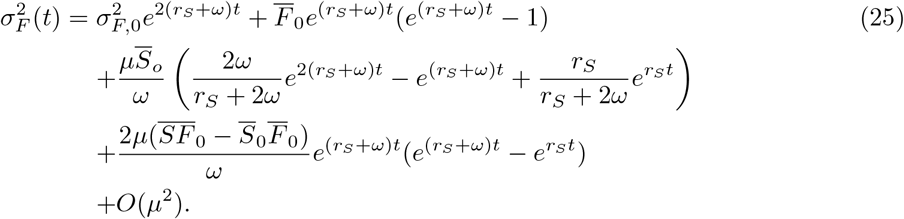

Using 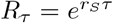 and *W*_*τ*_ = *e*^*ωτ*^, we rewrite

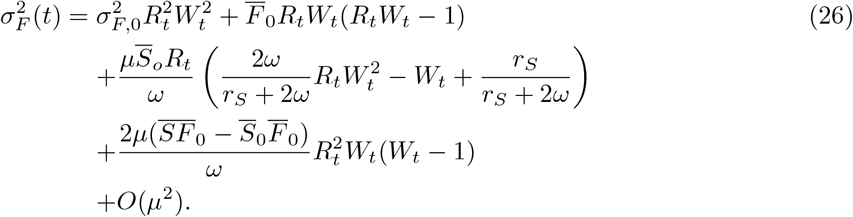

Using Eqs. (10),(11),(20),(25), we construct pdfs for *S*(*t*) and *F* (*t*) at the end of cycle *t* = *τ*. Then, we randomly sample a number from *P* (*S, τ*) for *S*(*τ*) and another number from *P* (*F, τ*) for *F* (*τ*). Those two numbers are cell numbers in a single Adult. We repeat this process for each Newborn to get cell numbers of all Adults. Note that the initial values for the Newborn *i* are 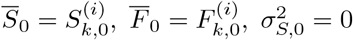,and 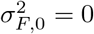. This process only requires two random numbers per collective while the result is consistent with the individual-based simulation.

**FIG. 1.**
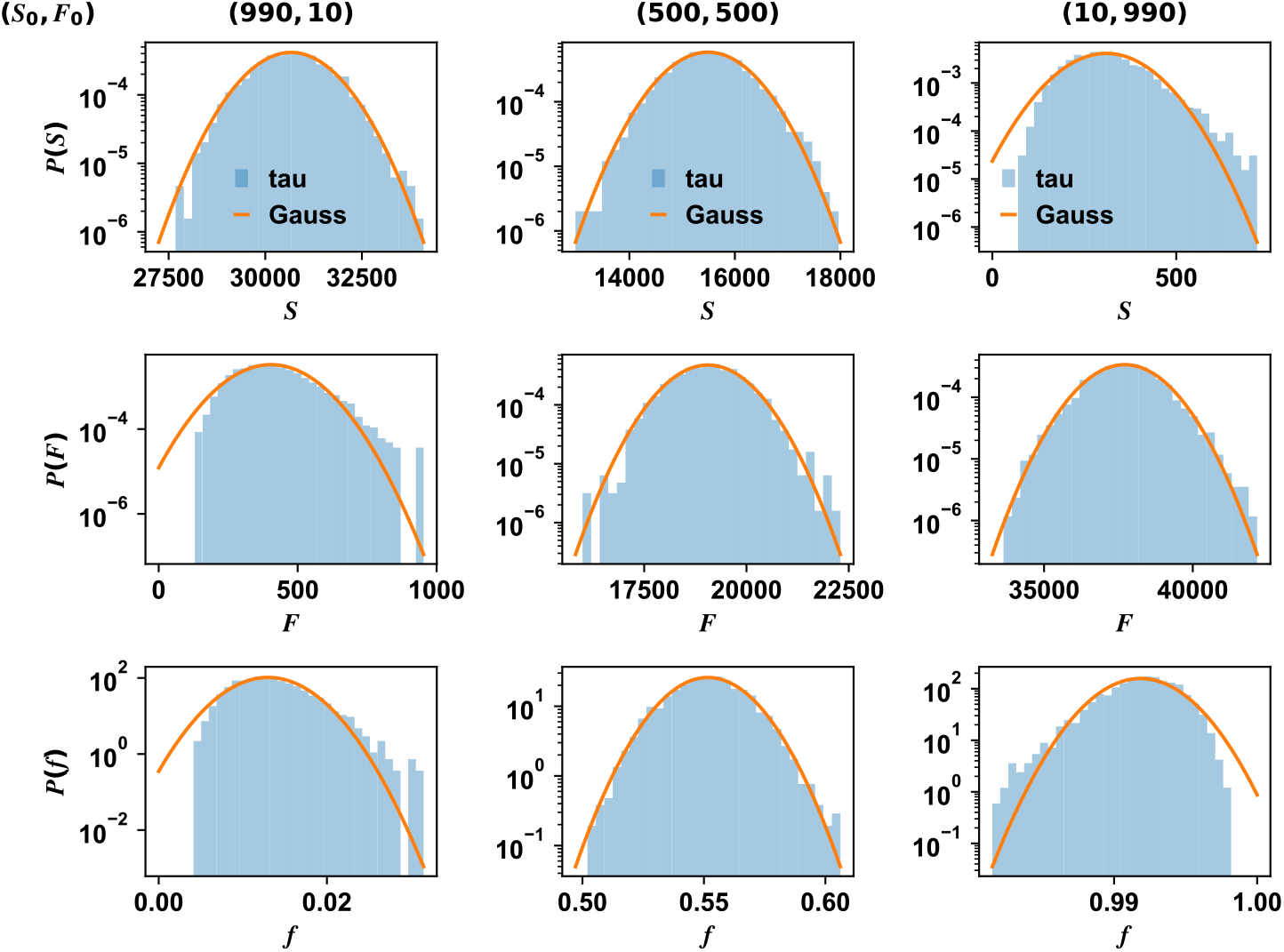
Comparison between the calculated Gaussian distribution (“Gauss”, with the mean and variances computed from Eqs. (10),(11),(20),(25)) and simulations using tau-leaping (“tau”). The simulations run 3000 times. The initial number of cells are (*S*_0_, *F*_0_) = (990, 10), (500, 500), and (10, 990) for each column. The parameters *r* = 0.5, *ω* = 0.03, *µ* = 0.0001, and *τ* = 4.8 are used.

Now, we check validity of Gaussian approximation for probability density functions of S and F populations. If we consider mutation from S to F as death in S population, then the process in S corresponds to a branching process with death. Also, the birth process in F including mutation results in a Luria-Delbrück distribution[3]. Thus the distributions of Adults’ S and F numbers are more skewed and heavy-tailed than Gaussian. But this problem is alleviated by larger initial S and F numbers and when the maturation time *τ* is not very long (see Fig. 1). Since we usually consider larger initial cell numbers, we use the Gaussian approximation on S and F populations in further calculations.

#### Selection

After sampling cell numbers of each Adult in the maturation step, we compute the F frequencies in each collectives 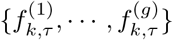.We denote the F frequency of collective *i* at time *t* = *τ* in cycle *k* as 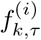.Among the *g* Adults, we select one collective with the F frequency which is the closest value to the target frequency 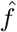.The selected Adult’s F frequency value is denoted by 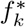.

**FIG. 2.**
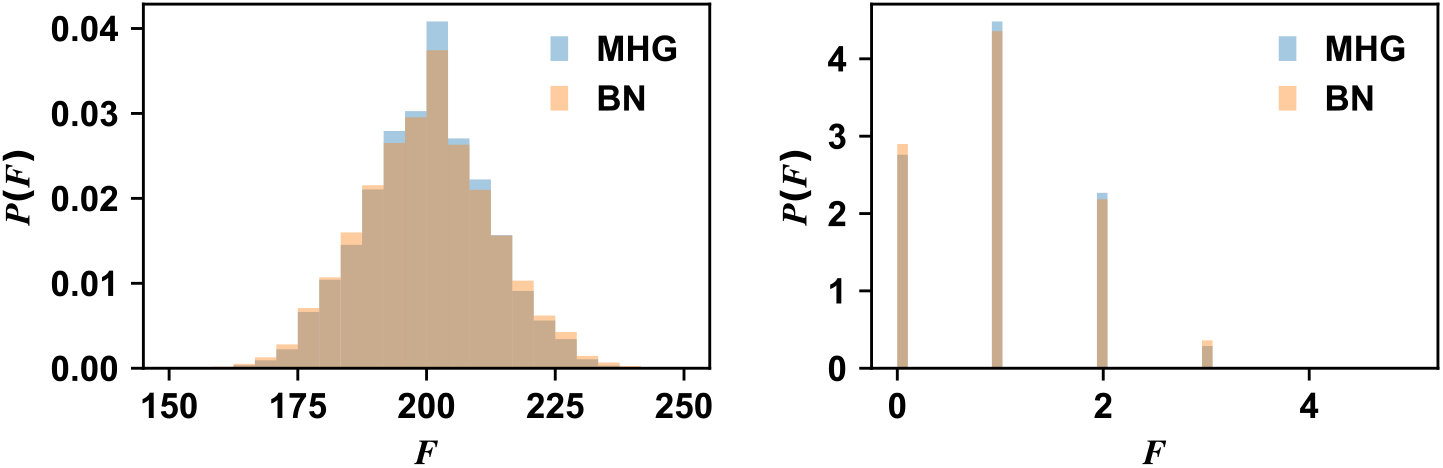
Congruence between consecutive sampling (MHG for multivariate hypergeometric distribution) and independent binomial (BN) sampling. The initial numbers of cells are *S* = 8000 and *F* = 2000 for the left panel, and *S* = 20 and *F* = 5 for the right panel. 10000 samples are drawn for each distribution. Here, a parent collective is divided into 10 collectives.

In mathematical expression, the selected frequency is defined by

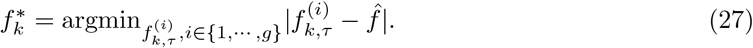

#### Reproduction

Using the chosen Adult, we generate *g* Newborn collectives for the next cycle *k* + 1. The most natural way is consecutive random sampling *N*_0_ cells from the selected Adult without replacement. In mathematical expression, we first randomly sample 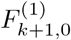 F cells and draw 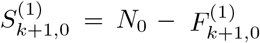 S cells from the selected Adult. Next, we sample 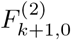 F cells and 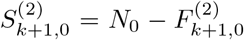 S cells from the remaining cells in the Adult. We repeat the process *g* times. Then the jpdf to choose 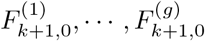 F cells, 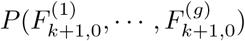, follows a multivariate hypergeometric distribution.

If we assume that the selected Adult size 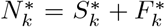 is large enough compared to Newborn size *N*_0_, the consecutive sampling is well approximated to the independent binomial sampling (see Fig. 2). Thus, we independently sample *g* numbers of F cells, 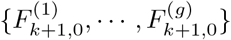, from the binomial distribution. The probability mass function of each 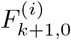 is given by

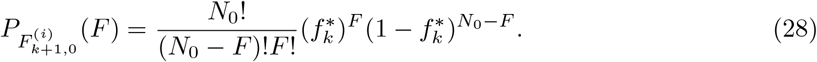

After sampling, the numbers of S cells are set to be 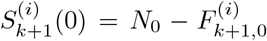 for each collective.

We can now start cycle *k* + 1 with these Newborn collectives. By repeating the above three steps (maturation, selection, and reproduction), we run the simulation until F frequency reaches a stationary state.

**FIG. 3.**
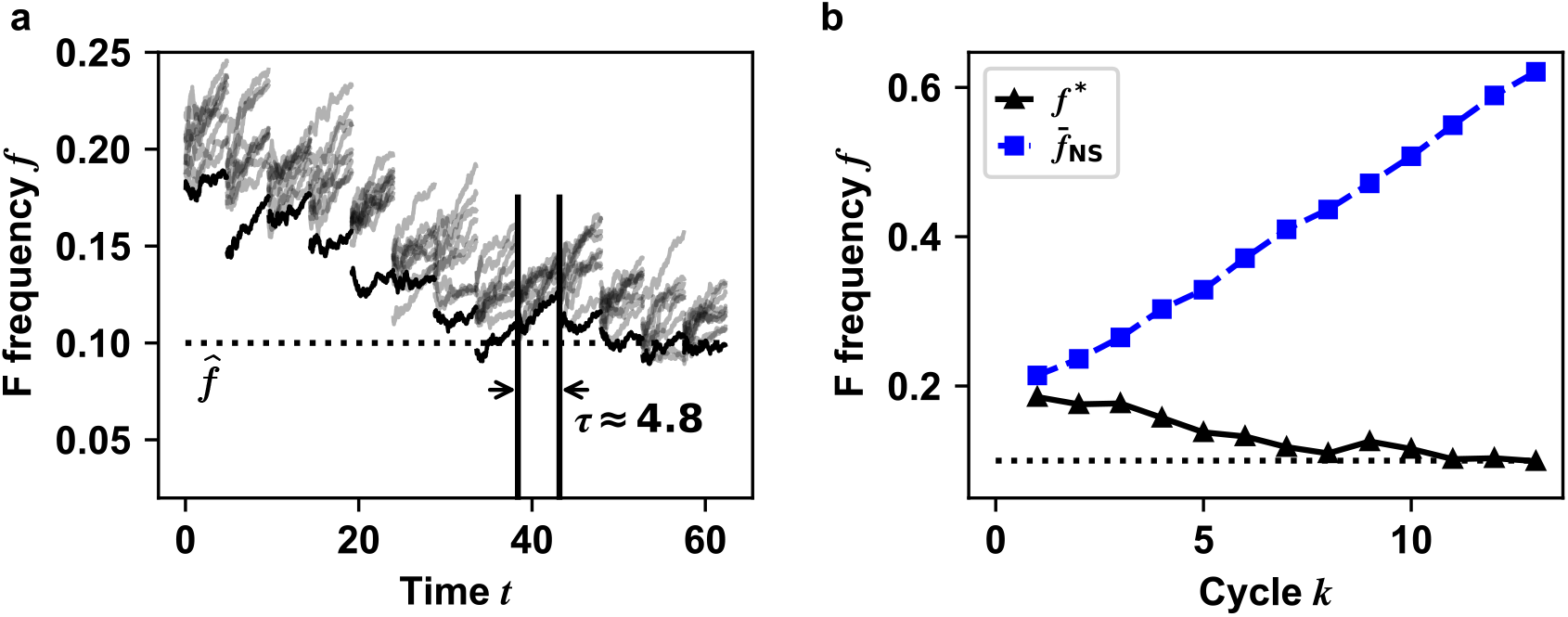
**a** Trajectories of F frequency for 10 collectives (*g* = 10) over time. The collective whose frequency is closest to the target value is selected in every cycle (black lines). The gray lines denote the other collectives. For parameters, we used S growth rate *r*_*S*_ = 0.5, F growth advantage *ω* = 0.03, mutation rate *µ* = 0.0001, maturation time *τ* ≈ 4.8, and *N*_0_ = 1000. **b** Comparison between frequency trajectories with selection (the chosen one Adult producing all offspring; black) and without selection (each Adult producing one offspring; blue) clearly shows the effect of artificial selection. The black line indicates F frequency of the selected collective 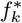 at each cycle in **a**. The blue line indicates the average trajectory without selection 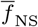 (the average of *g* = 10 individual lineages without inter-collective selection at the end of each cycle).

#### Simulation Result

Figure 3 presents the composition trajectories of all collectives using tau-leaping algorithm in the maturation step. The selected Adults have the closest composition to the target composition 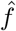.The selected Adult can have smaller F frequency than its parent Adult, so F frequency can be lowered after cycles.

In Fig. 4, we plot the absolute error *d* between the target frequency 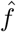 and 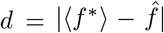 at the end of simulations (1000 cycles). Since the computing time for Tau-leaping algorithm (individual-based simulation) to reach 1000 cycles is very long, we used the sampling scheme in the above subsection. In the colormap, errors higher than 0.15 are marked with gray, which indicates selection failure. The dashed lines indicate the same boundary in Fig 2**e** in the main text.

### II CONDITIONAL PROBABILITY DISTRIBUTION OF THE SELECTED COLLECTIVE FREQUENCY *f* ^*^ AND SELECTION PROGRESS Δ*f*

In the main text, we identify the region of success by using selection progress 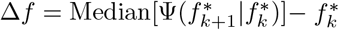,which is obtained from conditional pdf of 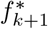 the F frequency of the selected Adult at cycle *k* + 1) given the selected 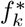 at cycle *k*, written as 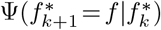.We consider the challenging case where 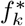 is above the target value 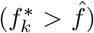,and therefore the Adult with minimum F frequency will be selected. To get an analytical expression of 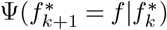,we first find the conditional pdf of *f* of Adults in cycle *k* + 1 given 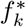 at cycle *k*. Then, we find 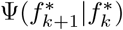 from the minimum value distribution of F frequencies among *g* Adults. Below, we describe mathematical details of this process.

**FIG. 4.**
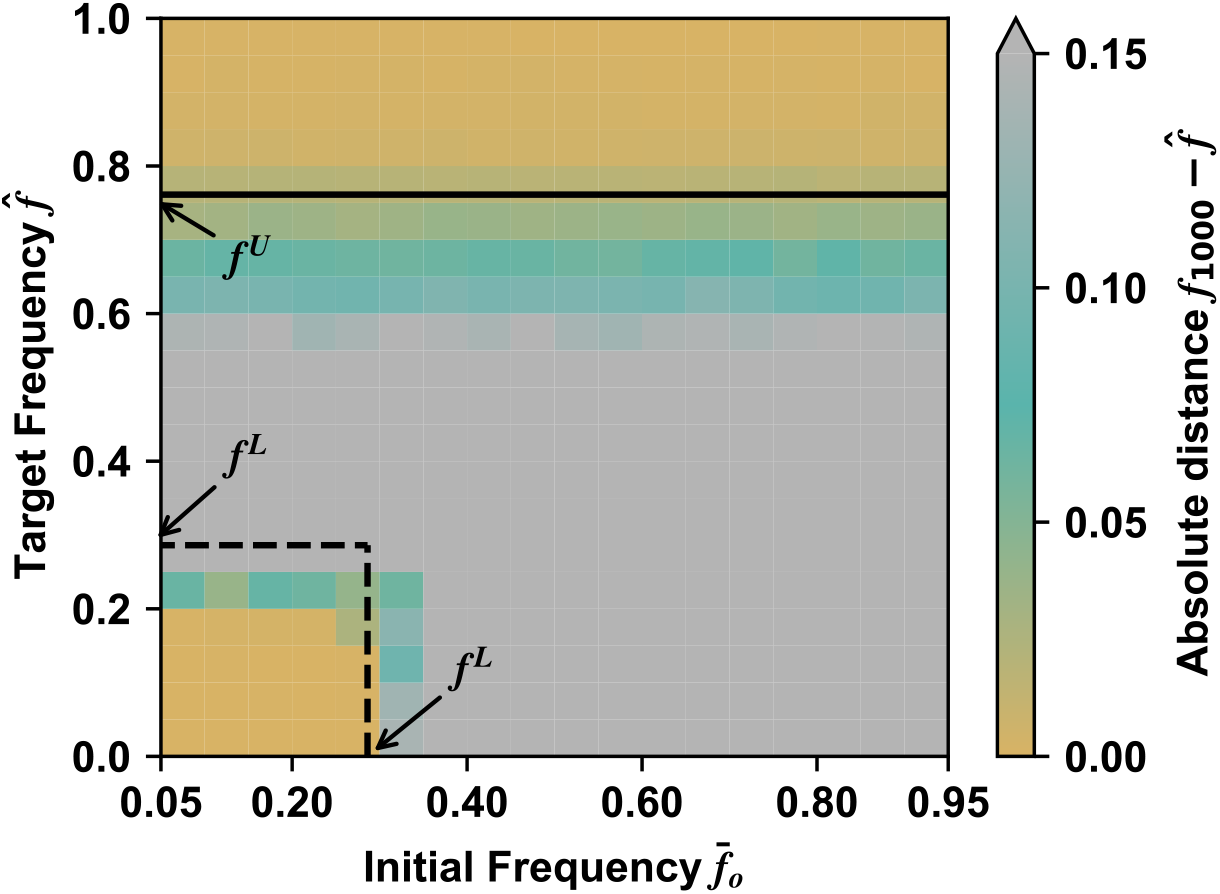
Color map of the absolute error 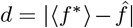 averaged selected collectives at the end of simulations (*k* = 1000) and the target frequency 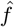. The solid and dashed lines are drawn by the arguments in the main text. For parameters, we used *r*_*S*_ = 0.5, *ω* = 0.03, *µ* = 0.0001, *N*_0_ = 1000, *g* = 10 and *τ* ≈ 4.8. The result is the average of 300 independent simulations. Compared to Fig. 2**e**, this figure has a higher resolution.

Let us start from the reproduction step from the selected Adult in cycle *k*. We reproduce *g* Newborns in next cycle *k* + 1. Then the probability distribution of the F cell numbers in Newborn collectives is given in Eq. (28). If the total number of cells in a Newborn collective *N*_0_ is large enough, Eq. (28) is approximated by the Gaussian distribution 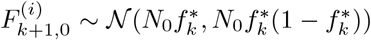. Then, the probability density function that 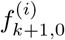 to be *ζ* in Newborn collective *i* is

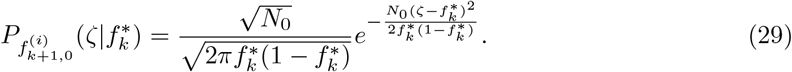

The Newborn collective *i* has initial cell numbers 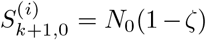 and 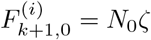.From here, we ignore cycle index *k* + 1 in subscript and *i* superscript for convenience.

Next, we write conditional pdf of Adults’ F frequency with given Newborn F frequency *ζ*. We assume that cell numbers in Adult *S*(*τ*) and *F* (*τ*) follow Gaussian distributions as in Eqs. (5) and (6). Based on Eqs. (10), (11), (20), and (25), we have

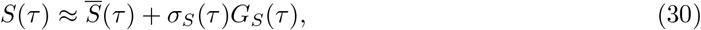

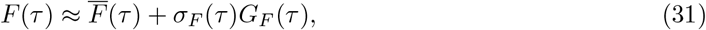

where *G*_*S*_(*τ*) and *G*_*F*_ (*τ*) are random variable following the standard distribution 𝒩 (0, 1). Note that each Gaussian is sharp if Newborn size *N*_0_ is sufficiently large (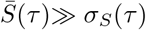and 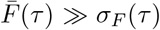. Then, we can approximately write *f* (*τ*) as

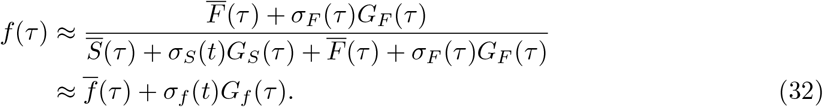

The mean of *f* is given by

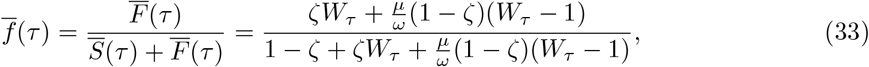

and the variance is

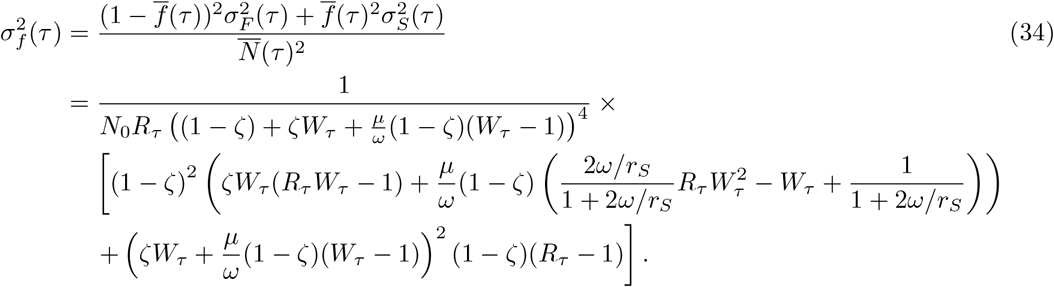

where 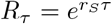 and *W*_*τ*_ = *e*^*ωτ*^. The average Adult size is 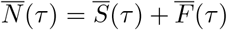.Thus the Adult’s F frequency *f* (*τ*) = *f*_*τ*_ follows the Gaussian distribution 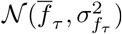 whose pdf is given by

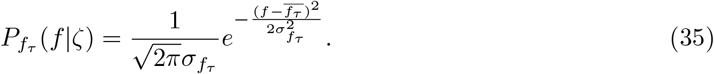

Next, we get the conditional pdf of 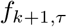 (offspring Adult’s F frequency in cycle *k* + 1) given 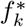.We multiply Eqs. (29) and (35) and take integral over *ζ*:

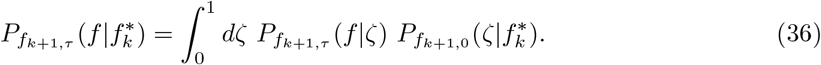

After maturation in cycle *k* + 1, the Adult with the smallest frequency is selected among *g* Adult collectives, denoted as 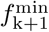.The pdf of 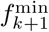 is obtained by the theory of extreme value statistics [4]. The cumulative distribution function (cdf) of the minimum value 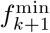 is given by

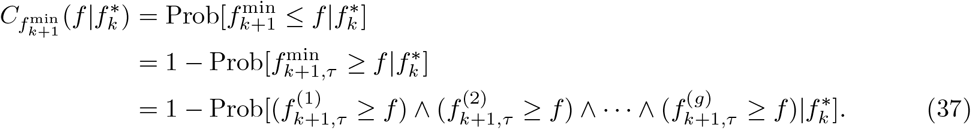

**FIG. 5.**
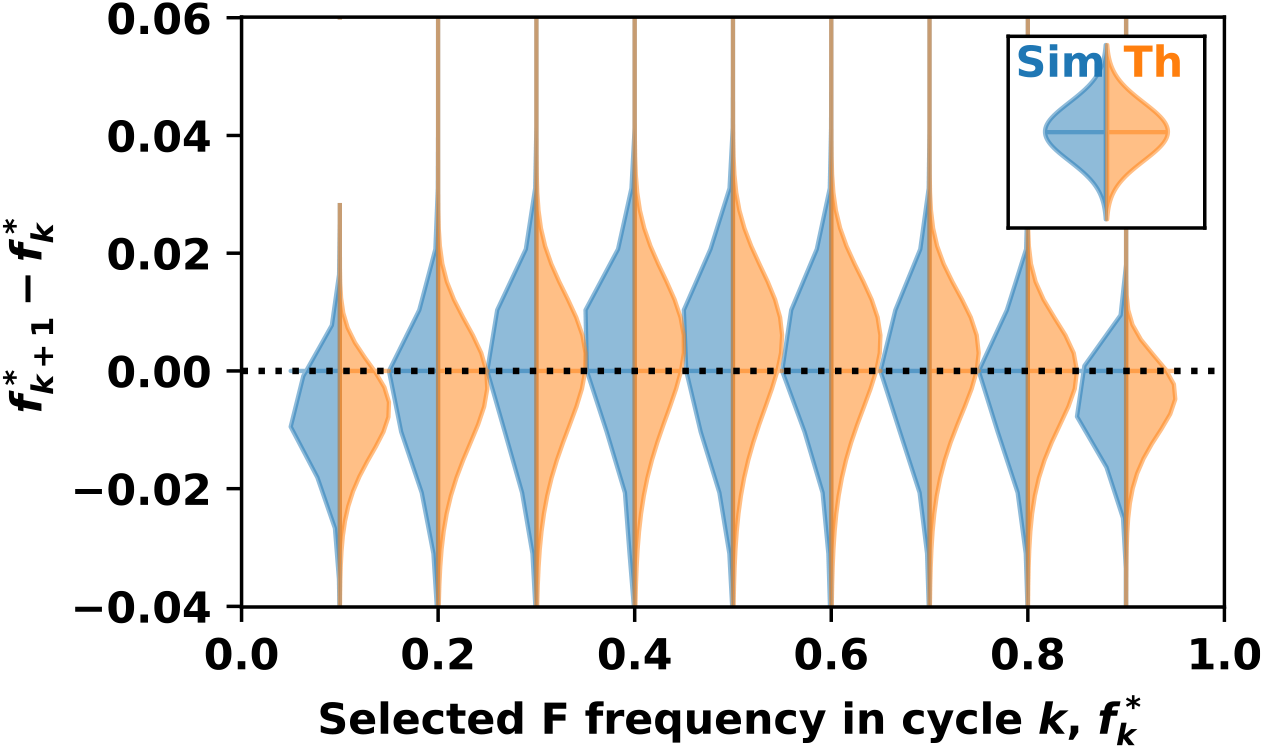
The probability density functions of the selected Adult’s F frequency 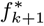 subtracted by 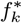. For simulations (blue), at each 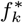,we performed 1000 stochastic simulations. The orange distribution represents Eq. (39) computed by numerical integration. The median values of the distributions are shown in Fig. 4**a** in the main text.

Since frequencies are independent and identically distributed, 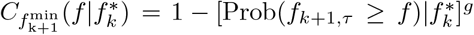. Note that Prob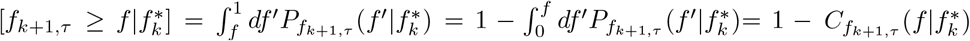, and Eq. (37) becomes

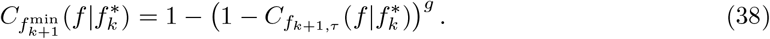

Then, the probability density function 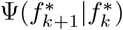 is obtained by differentiating Eq. (38) with respect to *f* and replacing 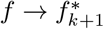,

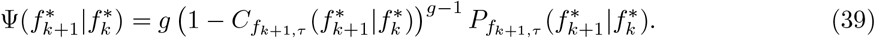

We compute the probability density function(39) by using numerical integration and compare it with the stochastic simulation results in Fig. 5. The two distributions are similar.

To get the analytic approximation of the median of Eq. (39), we assume that the Adult’s F frequency distribution is Gaussian. Then we only need to calculate the mean 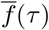 and variance 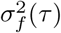 of Adult’s F frequency. Instead of calculating the integral with respect to *ζ* in Eq. (36), we put a set of initial value from Newborn’s F frequency distribution 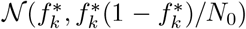 in Eqs. (12),(13),(20),(26): 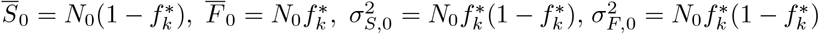,and 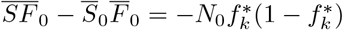. Then we have

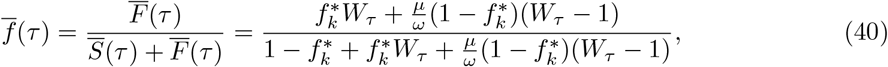

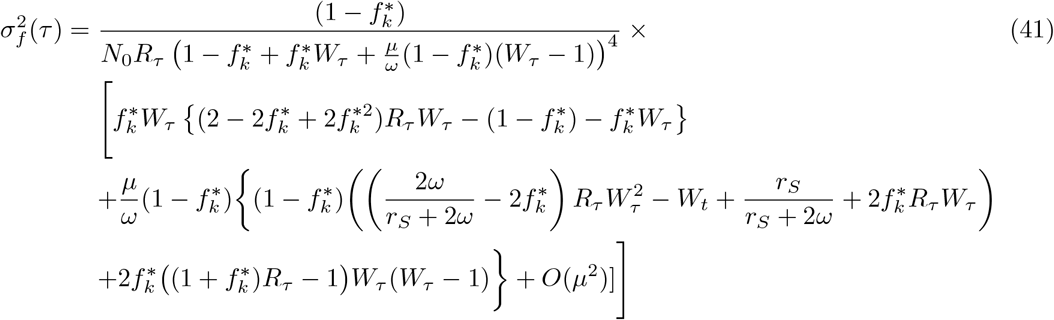

which give rise to Eq. (3) and Eq. (4) in the main text, respectively. The functional form of Eqs.(40) and (41) are plotted in Fig. 6**a**.

**FIG. 6.**
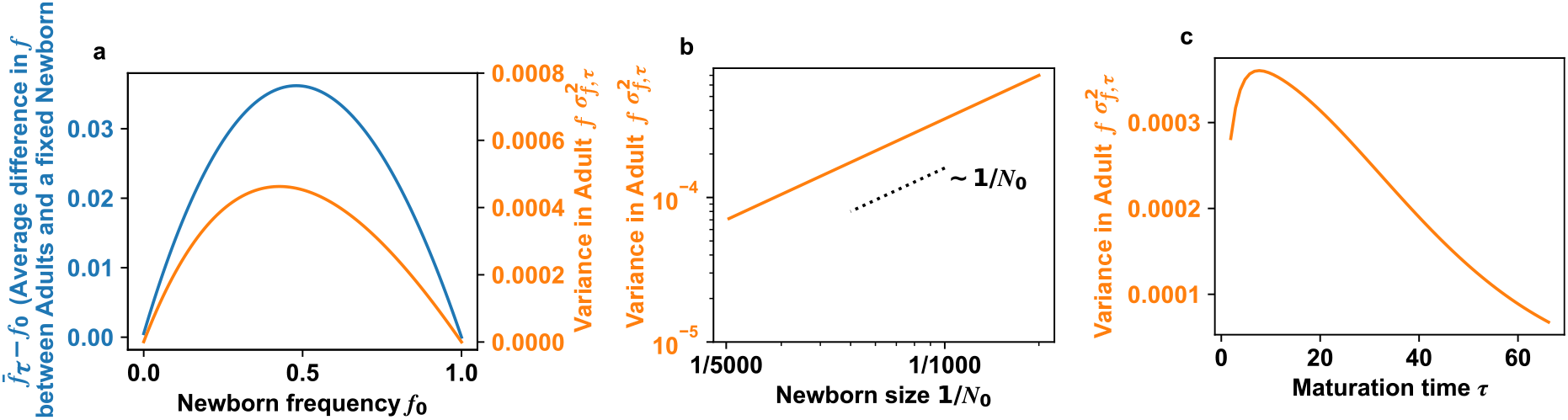
**a** Mean (Eq. (33)) and variance (Eq. (34)) of *f* values of Adult collectives with respect to the Newborn frequency *f*_0_. **b** Scaling relation of F frequency variance (Eq. (41)) with Newborn collective size *N*_0_. The initial F frequency is 0.5. The parameters are *r*_*S*_ = 0.5, *ω* = 0.03, *µ* = 0.0001, and *τ* ≈ 4.8. **c** Relation of F frequency variance (Eq. (41)) with maturation time *τ*. Other parameters are the same as **b**.

The median (Median 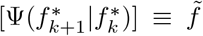) of Eq. (39) satisfies 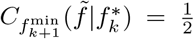,which means 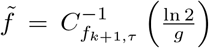. If we assume that the distribution Eq. (39) is Gaussian, then the inverse function 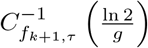 can be written as

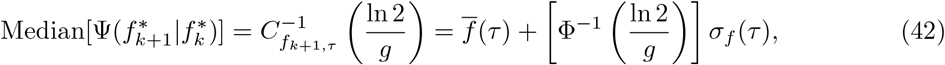

where Φ^−1^(*y*) is an inverse cumulative density function (CDF) of the normal distribution with mean 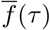 in Eq. (41) and standard deviation *σ*_*f*_ (*τ*), a square root of Eq. (41). Subtracting 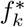 from Eq. (42) gives the selection progress

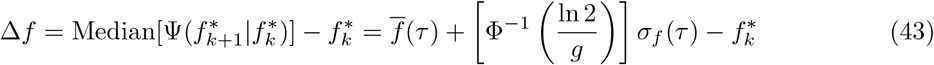

which is Eq. (2) in the main text.

**FIG. 7.**
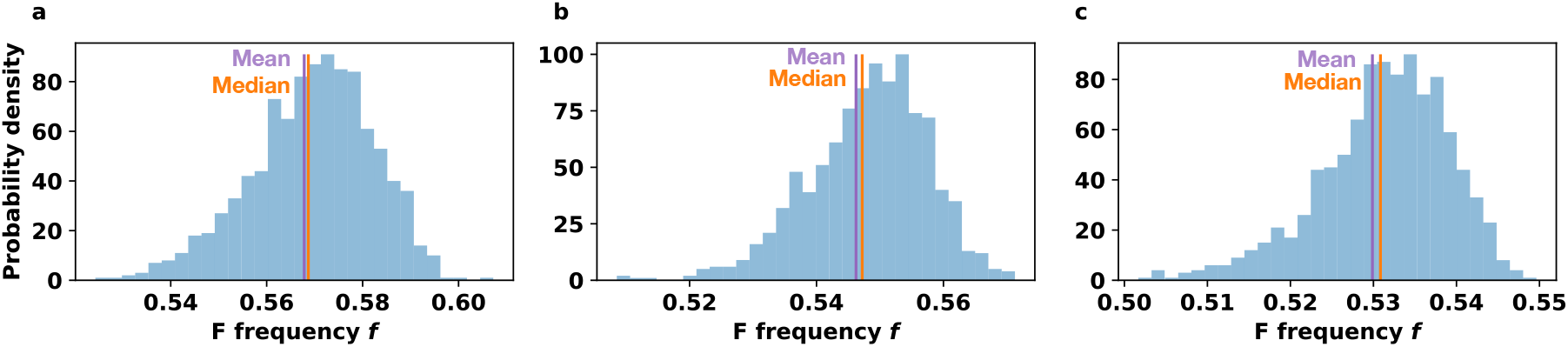
Median (orange) and mean (violet) have similar distributions. We performed 1000 simulations to get probability density. **a** *g* = 10, **b** *g* = 100, and **c** *g* = 1000. Initial F frequency is 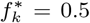.The parameters are *r*_*S*_ = 0.5, *ω* = 0.03, *µ* = 0.0001 and *τ* = ln[1000]*/r*_*S*_.

Further, we get an asymptotic expression of Φ^−1^(ln 2*/g*) when *g* is large (or Φ^−1^(*y*) with small *y*).

Here, we introduce a method from [5]. We start from the CDF of the standard normal distribution, 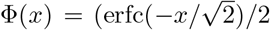 where the function erfc(*x*) is the complementary error function. To get the expression of Φ^−1^, we need an asymptotic expression of inverse of *y* = erfc(x) function (*x* = erfc^−1^(*y*)) as the inverse CDF 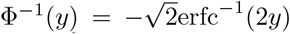.The known asymptotic expansion of *y* = erfc(*x*) for large *x* is 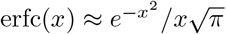.By taking logarithm of both sides, we have

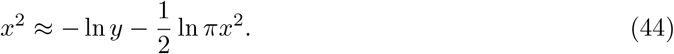

Replacing *x*^2^ on the righthand side in Eq. (44) into the expression itself, we get continued logarithmic form of

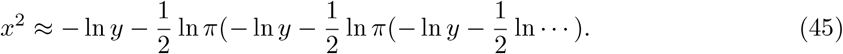

Inserting *x* = erfc^−1^(*y*) (square root of Eq. (45)) into the inverse CDF 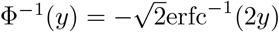,we have 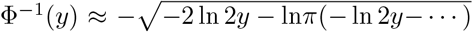.So, the asymptotic expression of Φ^−^ (ln 2*/g*) is given by

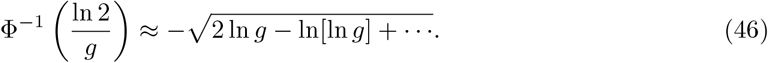

### III CRITICAL NEWBORN SIZE 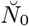 TO ALLOW ALL TARGET FREQUENCIES

First we note that 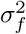 in Eq. (41) is proportional to 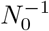 for the following reasons. Variance 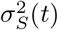 in Eq. (20) scales linearly with *N*_0_ since both 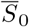 and 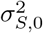 scale linearly with *N*_0_. Variance 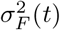 in Eq. (25) also scales linearly with *N*_0_ because 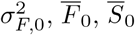,and covariance 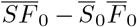 all scale linearly with *N*_0_. The mean Adult size 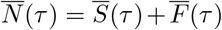 is also proportional to *N*_0_ because the average cell numbers in Eqs. (10) and (11) are linear with respect to *N*_0_. Thus, the scaling relation of Eq. (41) is given by 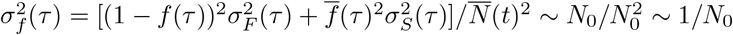.

Small *N*_0_ makes all target frequencies achievable shown in Fig. 4**a** in the main text. That is because small *N*_0_ induces large *σ*_*f*_, and thus *N*_0_ smaller than a certain critical value 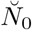 makes the selection progress Δ*f* always negative regardless of the value of 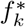 (*i*.*e*.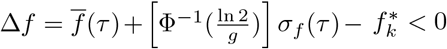). That means the inter-collective selection overcomes intra-collective selection in any target frequencies. To get an analytical approximation of the critical newborn size 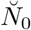,we simply assume that selection progress Δ*f* is maximum at 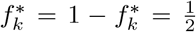 where the changes in 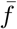 and 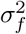 are fastest. If the maximum value of Δ*f* is zero, all other values of Eq. (42) are negative, which naturally states all targets are achievable. Putting 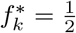,Eqs. (40) and (41) become

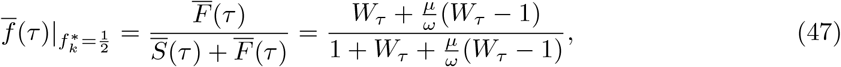

and

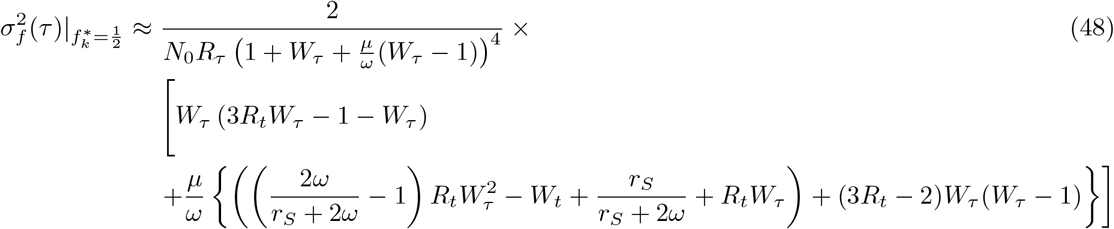

So, by setting 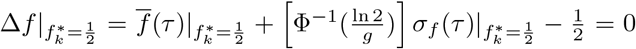 with 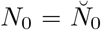,we get a solution of

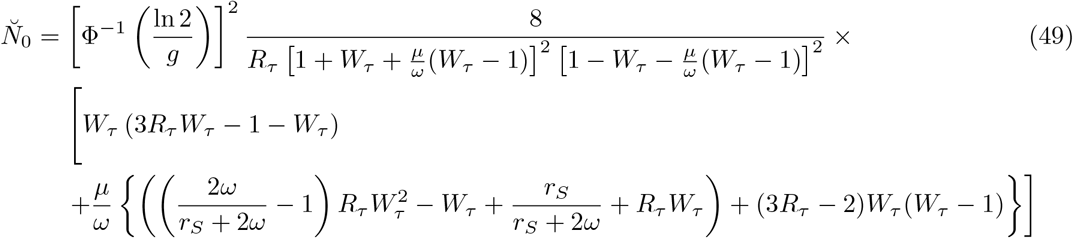

Thus, all target frequencies are successfully selected with Newborn size *N*_0_ smaller than 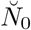. If mutation rate is zero, the critical value becomes

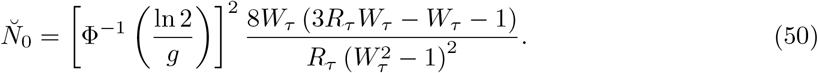

### IV SELECTION WITHOUT MUTATION *µ* = 0

When the mutation rate is zero, two genotypes behave as two distinct species. The compositional change is provided by Eq. (42) with setting *µ* = 0. Corresponding 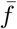 in Eq. (40) and 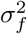 in Eq.(41) become

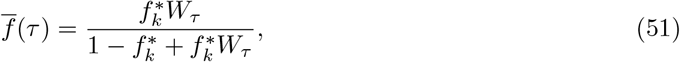

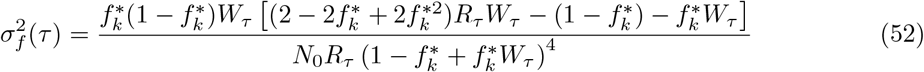

**FIG. 8.**
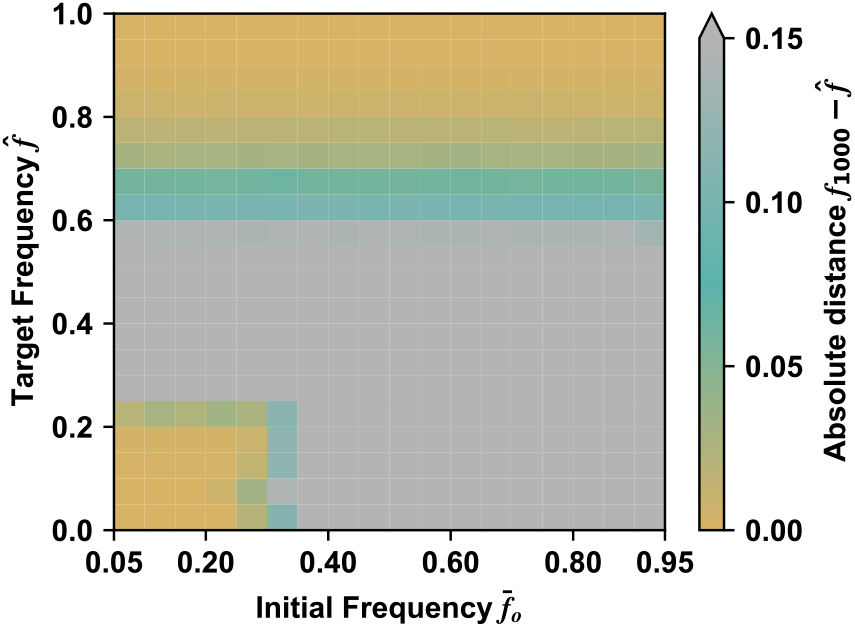
Simulation with zero mutation rate. Color map of the absolute error 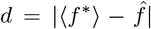 between frequency ⟨*f*\*⟩ of the averaged selected collectives at the end of simulations (*k* = 1000) and the target frequency 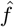.For parameters, we used *r*_*S*_ = 0.5, *ω* = 0.03, *µ* = 0, *N*_0_ = 1000, *g* = 10 and *τ* ≈ 4.8.

Equations (51) and (52) suggest that when a community consists of two competing species, we obtain similar conclusions on the accessible region for target composition. The stochastic simulation results are present in Fig. 8.

### V. STRONGER OR WEAKER ADVANTAGES *ω*

The solution of equation (2) in main text provides the boundary values with varying the *ω*, the fitness advantage of F over S. We numerically calculate the solutions and plot in Fig. 9.

### VI. DELETERIOUS MUTATION *ω <* 0

In the main text, we show the target composition can be achieved in some ranges of initial and target values when the mutation is beneficial to growth. The same analogy can be applied when the mutation is deleterious. Since the F cells grow slower than the S cells (*ω <* 0), the F frequency naturally decreases in the maturation step. Then, the challenging case is selecting larger F frequency against to the intra-collective selection. So the conditional probability distribution 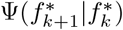 that we consider now is a maximum value distribution of Eq. (36). Thus, instead of Eq. (37), we look for the cumulative distribution function of the maximum value 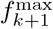 such that

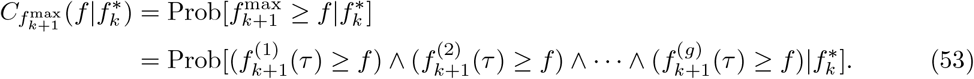

**FIG. 9.**
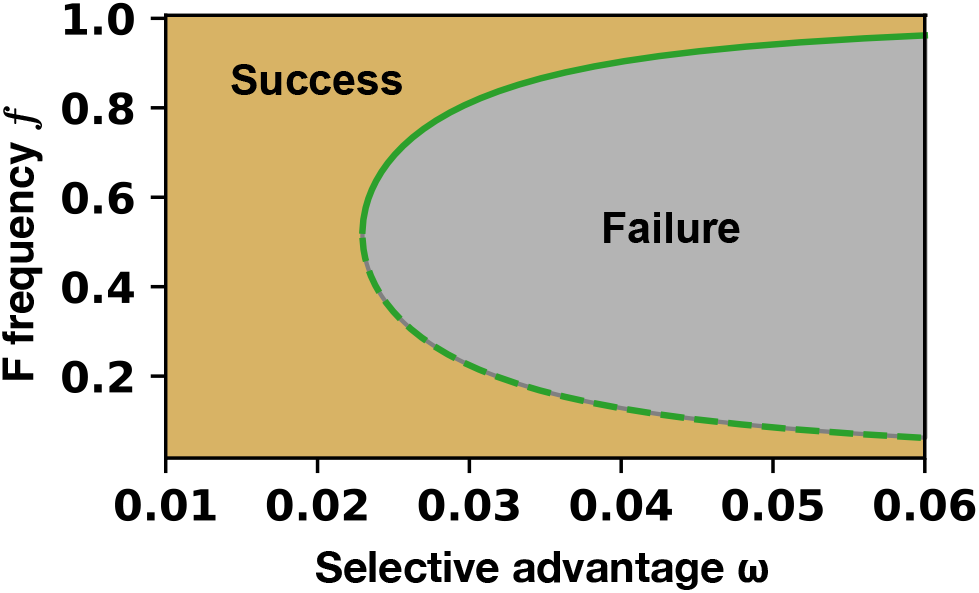
Change of success region in varying selective advantage *ω. r*_*S*_ = 0.5, *ω* = 0.03, *µ* = 0.0001, *N*_0_ = 1000, *g* = 10 and *τ* ≈ 4.8.

If all frequencies are independent and identical distributed random variables, the cumulative distribution function becomes

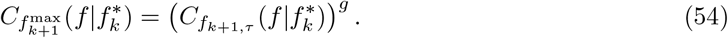

Likewise in the previous section, we get the conditional probability density function by differentiating Eq. (54) with respect to *f* and replacing 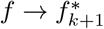 as

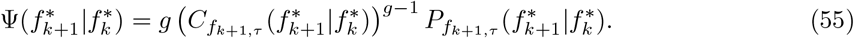

The distribution in Eq. (55) is evaluated for various 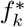 in Fig. 10**a** with numerical simulations, and the median values of distributions are presented in Fig. 10**b**. In the case of *ω* = −0.03, the target frequency lower than around 0.3 and larger than around 0.7 can be selected. Since the sign of *ω* is opposite to the result in the main text, the diagram is reversed from Fig. 2**e** in the main text.

### VII. SELECTING MORE THAN ONE COLLECTIVE

In the main text, we choose one collective which has the closest frequency to the target among *g* collectives. Such ‘top 1’ strategy allows us to apply extreme value theory. However, ‘top 1’ may be too restrictive [6]. Thus, we test the ‘top-tier’ strategy by choosing top 5 among 100 Adults (Fig. 11). The top-tier strategy is shown to be inefficient in our system. This is because in [6], non-heritable variations−such as stochastic fluctuations in species composition introduced by pipetting − caused nonheritable variations in collective function. Nonheritable variations could potentially mask desired mutations if these mutations happened to occur in an ‘unlucky’ environment that yielded lower collective functions. Hence, lenient selection would allow the preservation of these mutations. In contrast here, stochastic fluctuations in genotype composition is heritable: a parent Newborn with lower F frequency *f* will tend to have offspring Newborns with lower *f* values. Hence, top-1 is more effective in this study.

**FIG. 10.**
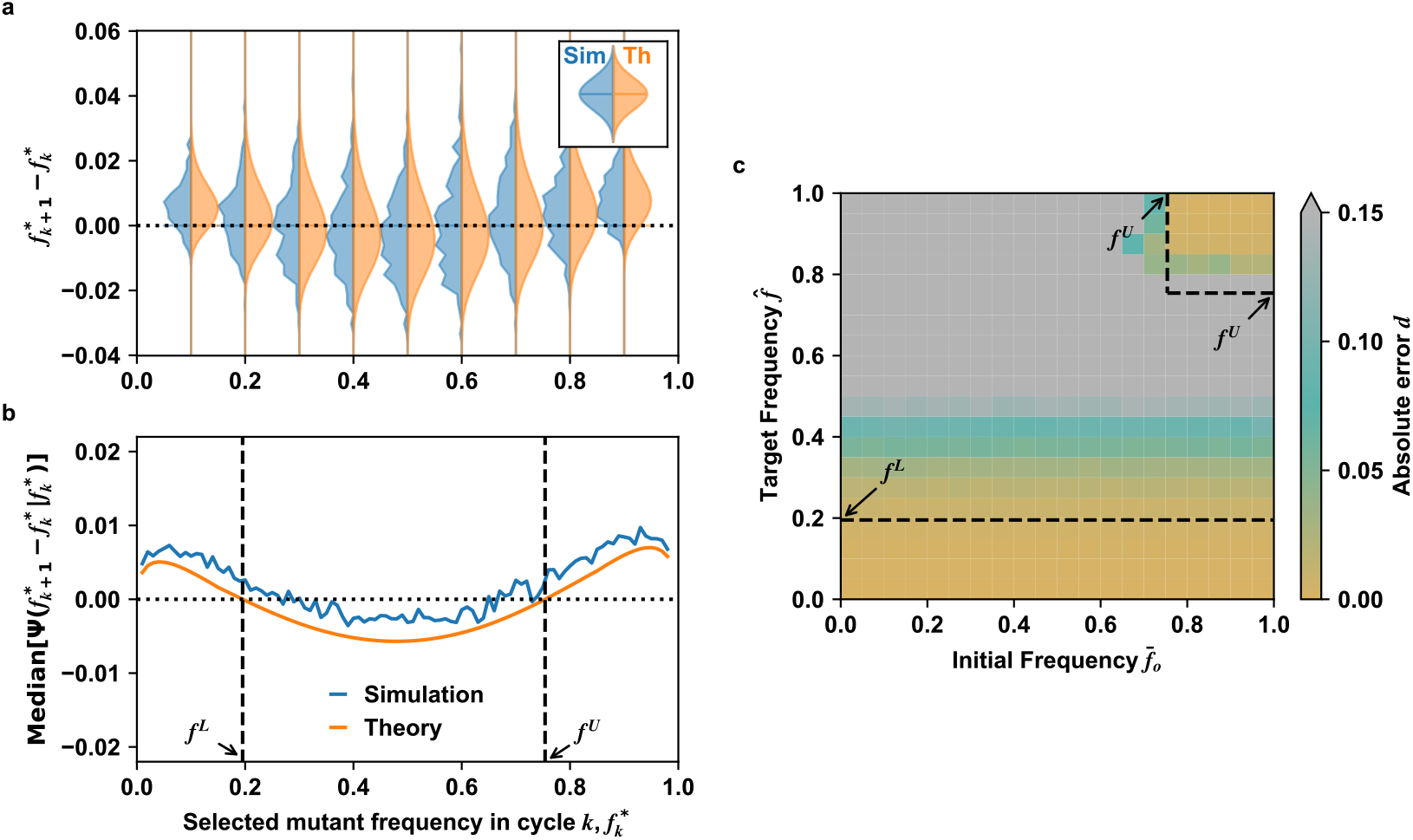
Artificial selection also works for deleterious mutation. **a** Conditional probability density functions of 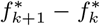 for various 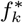 values. The left-hand side distribution is obtained from simulations and the right-hand side distribution is numerically obtained by evaluating Eq. (55). Small triangles inside indicate the median values of the distributions. **b** the median value of distributions at a given 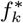.The points where the shifted median becomes zero, Median 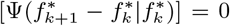 are denoted as *f* ^*L*^ and *f* ^*U*^, respectively. **c** The relative error between the target frequency 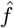 and the ensemble averaged selected frequency 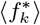 is measured after 1000 cycles starting from the initial frequency 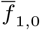.Either the lower target frequencies or the higher target frequencies starting from the high initial frequencies can be achieved. The black dashed lines indicate the predicted boundary values *f* ^*U*^ and *f* ^*L*^ in **a**.

### VIII. EXTENSION TO THREE-POPULATION SYSTEM

We assume that collectives consist of three genotypes with slow-growing(S), fast-growing(F), and faster-growing(FF) types. The growth rate of S is *r*_*S*_. Each mutation adds *ω* to the growth rate. Thus, the F and FF types have growth rates *r*_*S*_ + *ω* and *r*_*S*_ + 2*ω*, respectively. The mutation rate is *µ*. So, the birth and mutation events are written by the chemical reactions:

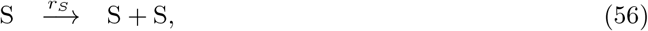

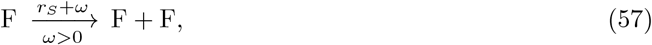

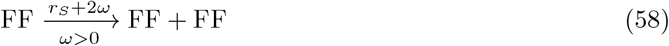

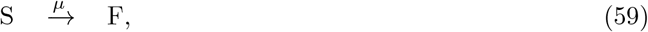

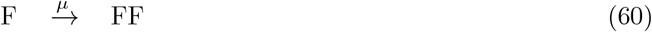

**FIG. 11.**
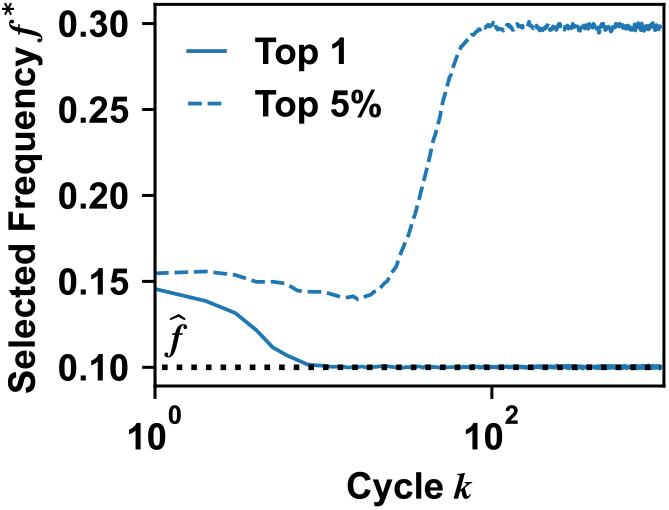
Selecting Top-5% outperforms selecting Top 1. We bred 100 collectives and chose either top-1 collective (solid line) or top-5 collectives (dashed line) with *f* closest to the target value 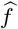 (black dotted line).

We write a master equation of the processes for *P* (*S, F, FF, t*) which is the probability to have *S, F* and *FF* numbers of S, F, and FF cells at time *t*, respectively.

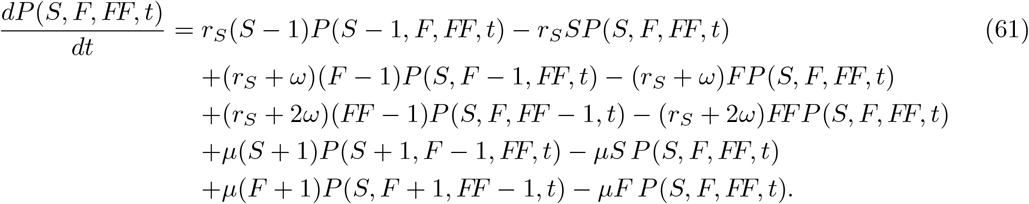

The composition of collective *i* in cycle *k* is now represented with two frequencies 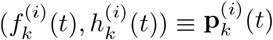 where the F frequency is *f* 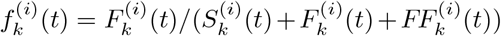 and the FF frequency is 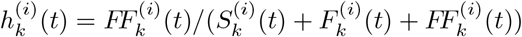.Then, the target composition is set to be 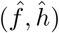. The composition of the selected Adult in cycle *k* is 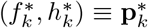.We apply the processes used in the above section II to obtain the conditional probability 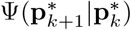 by using the master Eq. (61). At the reproduction step in cycle *k*, we choose *N*_0_ cells from the selected Adult whose composition is 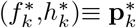.Then, Newborn collectives are independently sampled from a multinomial distribution. For convenience, we drop the collective index (*i*). Then, the conditional joint probability mass function of *F*_*k*+1,0_, *FF*_*k*+1,0_ cells is represented by

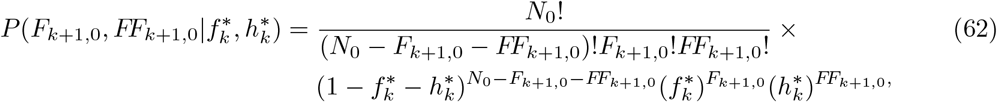

where the number of S *S*_*k*+1,0_ is automatically set to be *S*_*k*+1,0_ = *N*_0_ − *F*_*k*+1,0_ − *FF*_*k*+1,0_. Then, the approximated multivariate normal distribution is 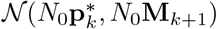 where the mean distribution is 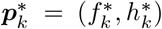 and covariance matrix is **M**_*k*+1_. The diagonal terms of **M**_*k*+1_ are variances 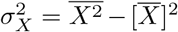 and the off-diagonal terms are covariances 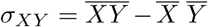.The matrix is given by

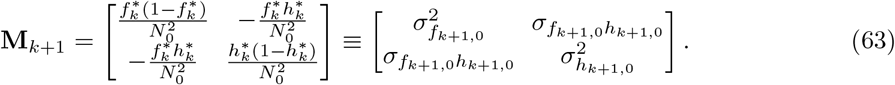

Then a Newborn’s composition ***ρ***(*ζ, η*) follows the multivariate Gaussian distribution 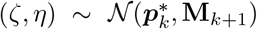 whose joint probability distribution is given by

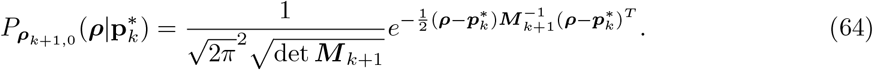

At the beginning of cycle *k*, a newborn collective starts from (*S*_0_, *F*_0_, *FF*_0_) cells (for convenience, cycle index *k* is dropped.) In terms of (*ζ, η*), each initial numbers are *S*_0_ = *N*_0_(1− *ζ* − *η*), *F*_0_ = *N*_0_*ζ*, and *FF*_0_ = *N*_0_*η*. Their initial covariance matrix is 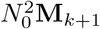.By using Eq. (61), we can write ordinary differential equations up to the second moment.

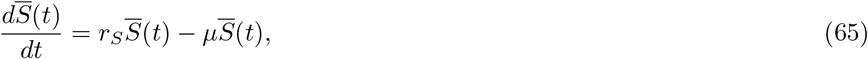

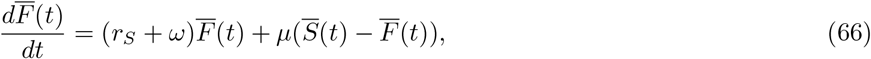

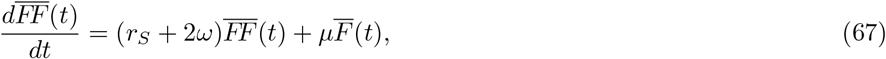

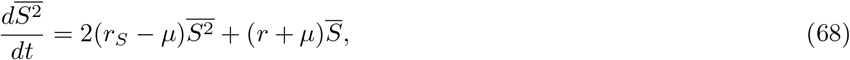

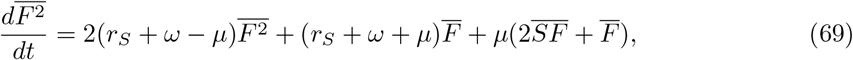

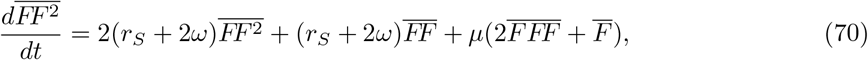

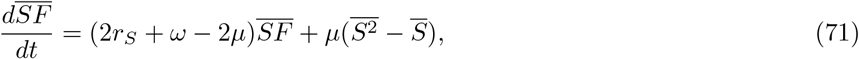

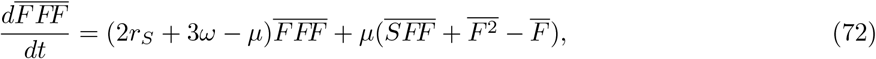

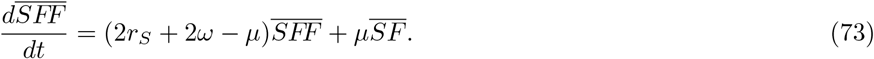

The initial conditions of the system in coupled Eqs. (65)-(73) are obtained by the mean and (co)variances of Eq. (62). By solving equations numerically, we obtain a set of mean cell numbers 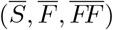 and a set of variances 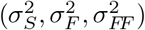 as well as covariances (*σ*_*SF*_, *σ*_*F FF*_, *σ*_*SFF*_). We assume that the covariances are smaller than the variances. We consider *S, F*, and *FF* as Gaussian random variables

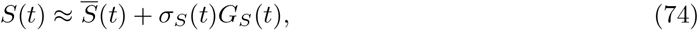

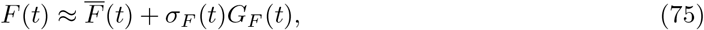

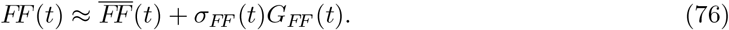

Then, the F frequency becomes

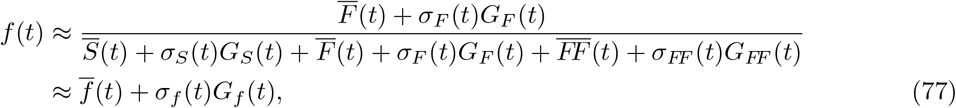

where 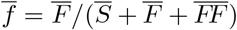 and 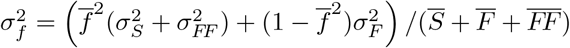. Similarly, the FF frequency is

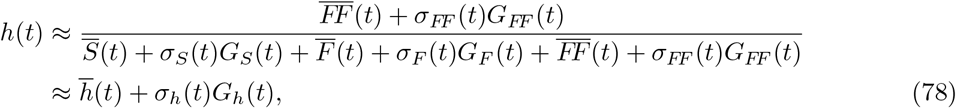

where 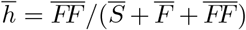 and 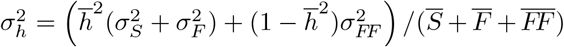.The dynamic flow of F and FF frequencies during maturation is shown in Fig. 12**a**. If the covariances are small enough, we can approximate the joint probability distribution of Adult’s composition (*f*_*τ*_, *h*_*τ*_) = **p**_*τ*_ as

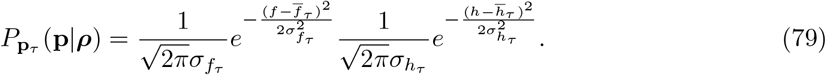

With cycle index *k*, we get conditional probability of matured collectives 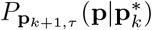 by

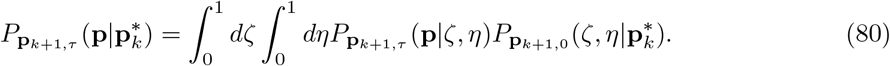

We select the Adult collective among *g* Adult collectives such that the change in frequencies during maturation could be compensated. During maturation, a frequency distribution moves in different directions in (*f, h*) space depending on the initial composition 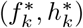 So, we take different directions to obtain the extreme value distributions. Considering only the sign of the frequency changes in *f* and *h*, we take either maximum or minimum. The mean change in *h* is always positive in the whole (*f, h*) space since 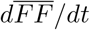 is always positive in Eq. (67). Thus, we choose the minimum value 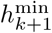 in every selection step.

If the mean 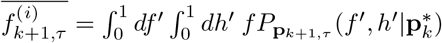 is larger than 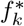,the minimum value among 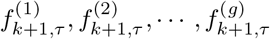 will be chosen in selection step to compensate the frequency change in maturation step. Let us denote the selected valued of *f* and *h* as 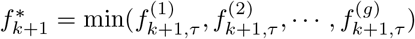 and 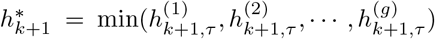. We temporarily drop time index *τ* for simplicity.

Then, the joint cumulative distribution function 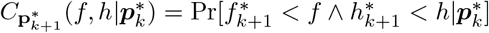 is

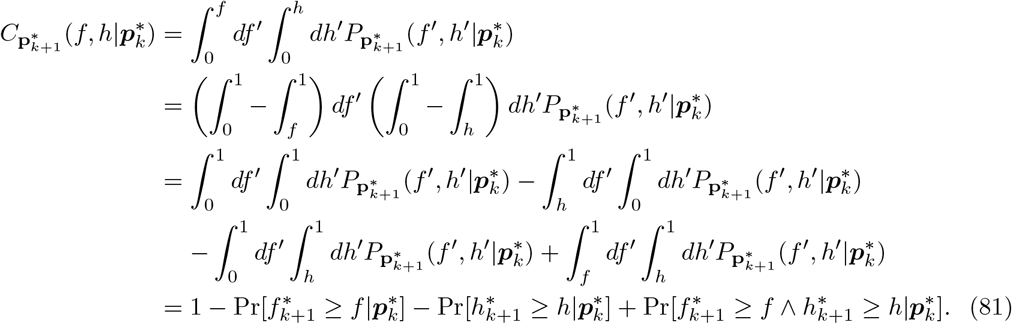

The probability 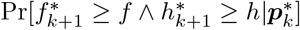 can be converted as

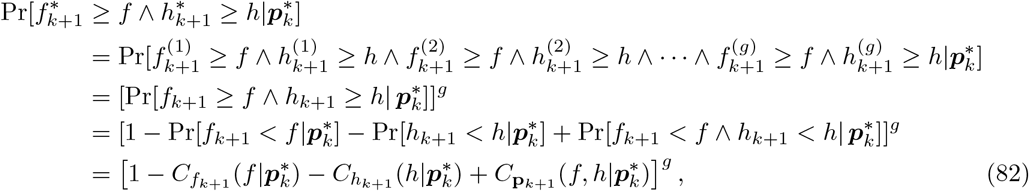

where 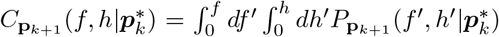 is a conditional joint cumulative distribution function of (*f* ^(*i*)^, *h*^(*i*)^). The marginal cumulative distribution functions are

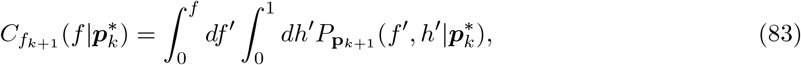

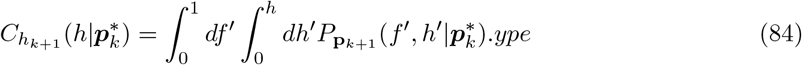

Similarly, the probabilities 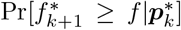 and 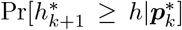 are converted into 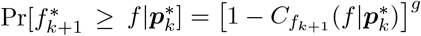 and 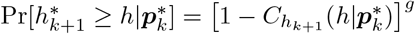. Thus, the joint cumulative distribution function is

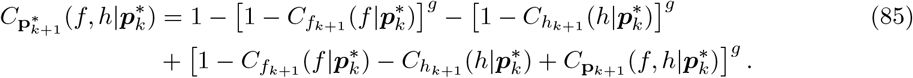

Then, the conditional probability of the selected collective is given by

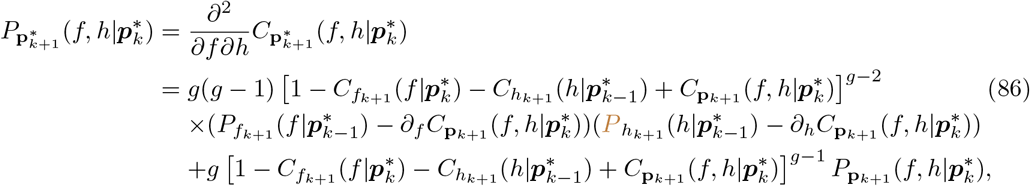

where 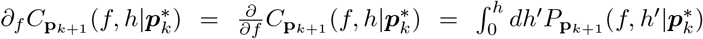 and 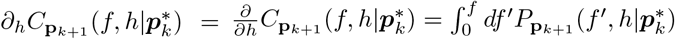

If the mean 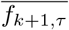 is smaller than 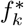,the chosen collective is likely to have maximum *f* values among *g* matured collectives. Then, the definition of *f* ^*^ is written by 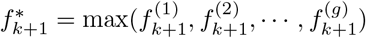.

We rewrite the joint cumulative distribution function 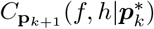 to be little different from Eq. (81) because now we have to utilize the condition *f* ^*^ *< f* instead of *f* ^*^ *> f*,

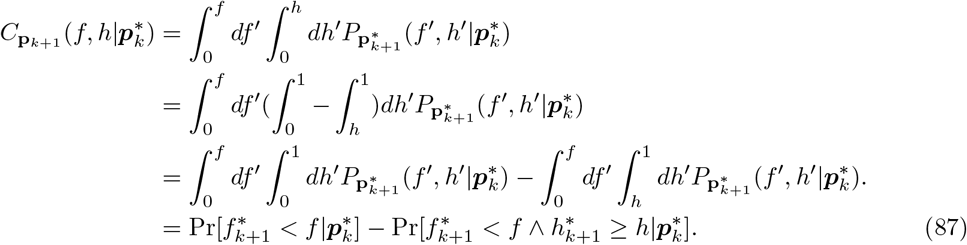

The probability 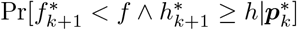 is converted as

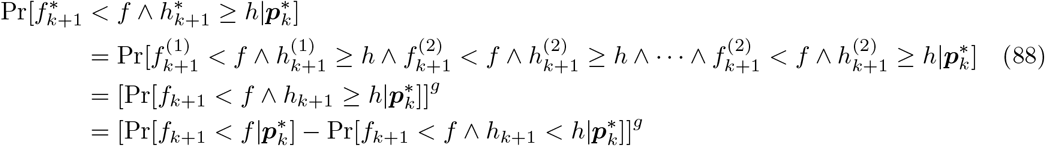

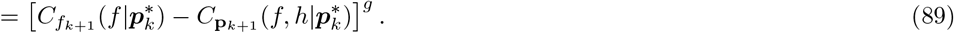

Thus, the joint cumulative distribution function is given by

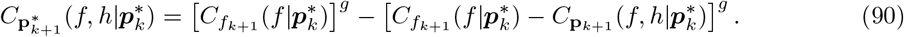

In this case, the conditional probability distribution function is given by

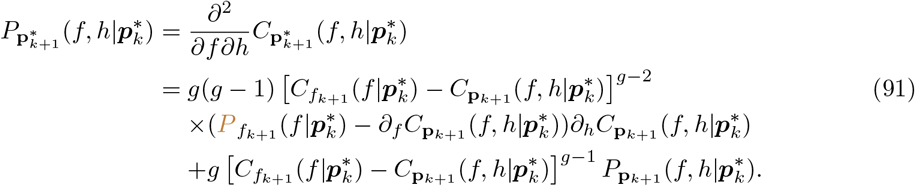

By replacing (*f, h*) to 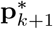,we finally obtain the conditional probability distribution 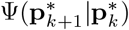,

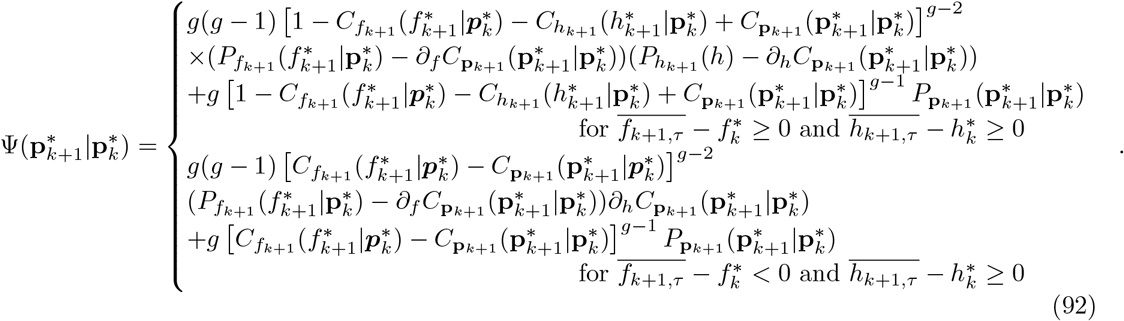

Using Eq. (92), we get the mean values of *f* ^*^ and *h*^*^ as

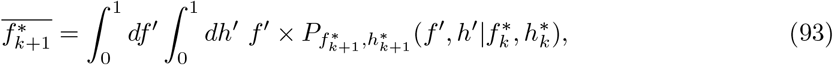

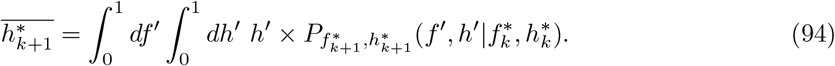

We define the accessible region in frequency space where the signs of the changes in both F frequency and FF frequency after a cycle are opposite to that of maturation (see Fig. 12),

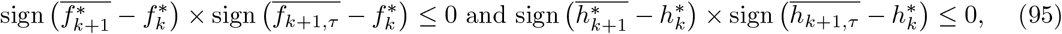

where 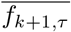 and 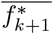 are the mean values of F frequencies after the maturation step in cycle *k* + 1 before and after selection, respectively, and *h* values are defined similarly for FF. Or, if the condition is not met, the composition of the selected collective may diverge from the target composition after several cycles. The accessible regions are marked in the gold-colored area in Fig. 12**b**. Similar to the two-population case, the accessible region is shaped by the flow velocity of the composition during the maturation step, as depicted in the flow diagram in Fig. 12**a**. Both F and FF frequencies tend to increase, and the inter-collective selection can compensate for these changes if the composition changes slowly when the F and FF frequencies are small. However, if the changes occur too rapidly when the FF frequency is intermediate, the frequency cannot be stabilized. So the accessible region is limited to the regions where the composition changes slowly.

This is explainable by projecting the three-population problem into the two-population problem. The selective advantage of FF relative to the rest of collective mainly determines the accessible region. The growth rate of the rest varies from *r*_*S*_ to *r*_*S*_ + *w* according to F frequency, so the mean growth rate of the rest is written by 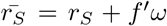 where *f* ^′^is F frequency in S+F. Then, the corresponding selective advantage of FF is 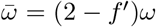 which varies between *ω* to 2*ω*. Using 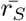 and 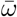 similar to Sec. II, we get bounds of accessible region (see dashed-line in Fig. 12**b**). The boundary from projected problem agreed well with the original three-population problem.

### IX DERIVATION OF EQUATIONS

In this section, we go over the derivation of Eq. (10)-(34) for readers not equipped with advanced mathematics training.

Assumptions: *µ ≪ ω, r*_*S*_

**FIG. 12.**
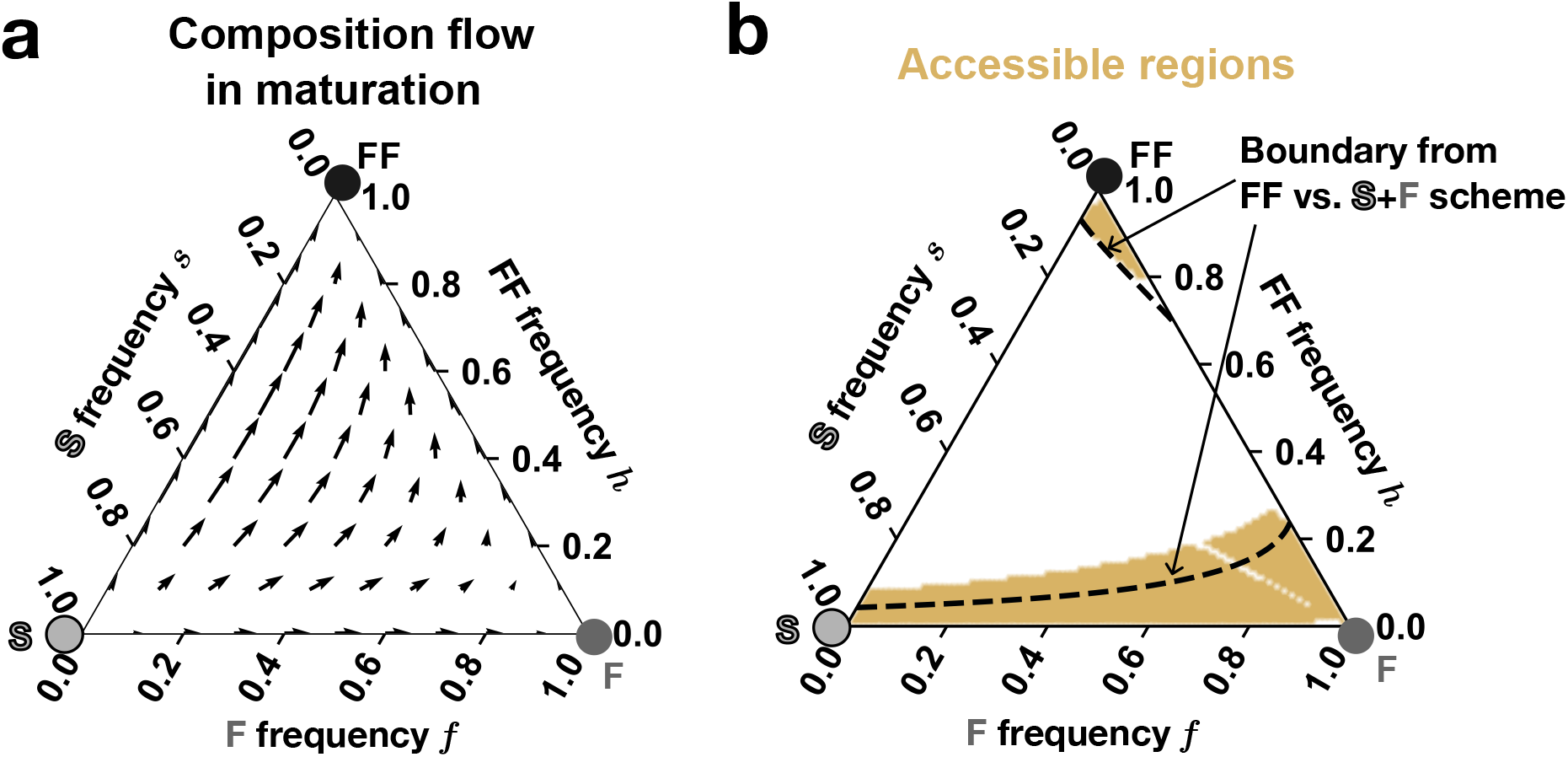
**a** The flow of composition change in F and FF frequencies at each composition (*f, h*). Top corner indicates that FF cells fix in the collective. Right bottom corner means collectives with only F cells while collectives contain S cells only at left bottom corner. Arrow length means the speed of change. **b** The accessible regions are marked by the gold area. If the signs of changes in both F frequency and FF frequency after inter-collective selection are opposite to those during maturation, then the given composition is accessible. Otherwise, the composition is not accessible and will change after cycles. Dashed lines are the boundary of accessible region by projecting the collective into a two-population problem (FF vs. S+F). The figures are drawn using *mpltern* package [7].

#### Equations (10) and (11)

Equation (10) is straightforwardly solved by integrating Eq. (8). Equation (11) is obtained from Eq. (9) using integration factor 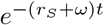:

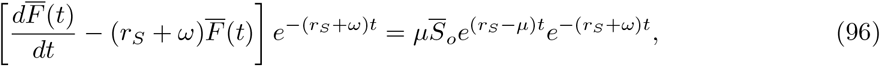

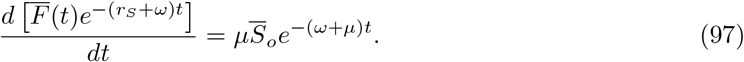

Integrating both sides, we get 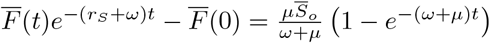.Thus,

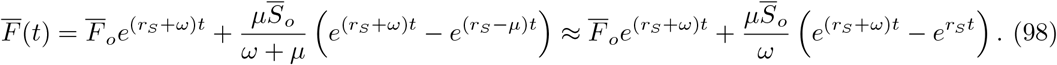

#### Equations (16) and (17)

Applying Eq. (4), we have

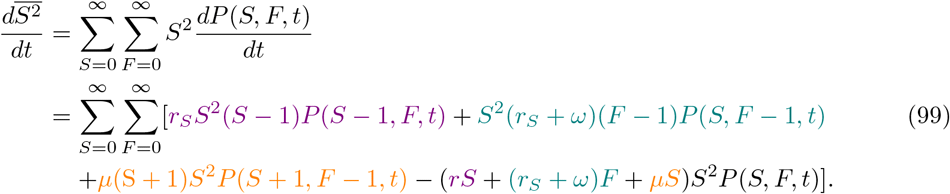

We collect the two violet-colored terms and change the order of summation. Note that the first violet-colored term does not change regardless of whether *S* starts from 0 or 1 because the term is zero for *S* = 0. Thus, the first violet-colored term is equivalent to 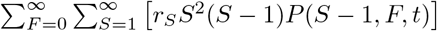. Let *α* = *S* − 1, and this becomes 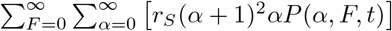. We reassign *α* as *S*, and obtain:

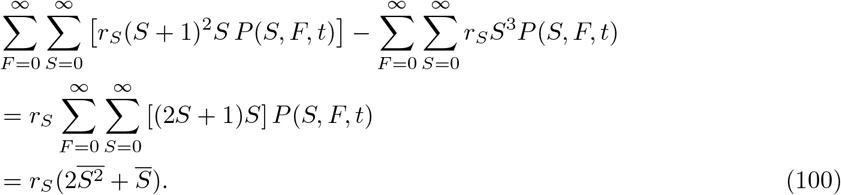

We collect the two green terms, and similarly obtain:

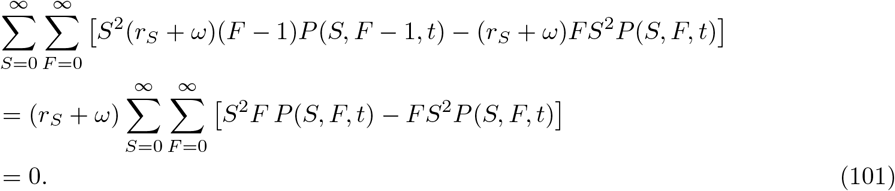

Finally, we collect the two orange terms. For the first orange term, the sum is the same regardless of whether we start from *S* = 0 or −1. Let *S* start from −1, and we have

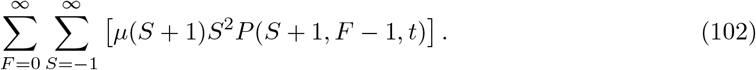

Let *α* = *S* + 1,then the term becomes 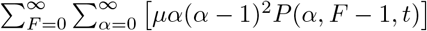.We reassign *α* as *S*, and additionally apply index change on *F* − 1:

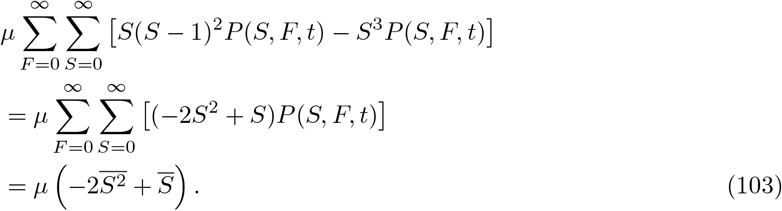

Now, add the three parts together, and we have

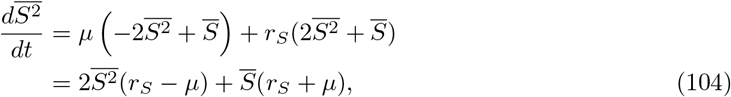

which is Eq. (16). Likewise,

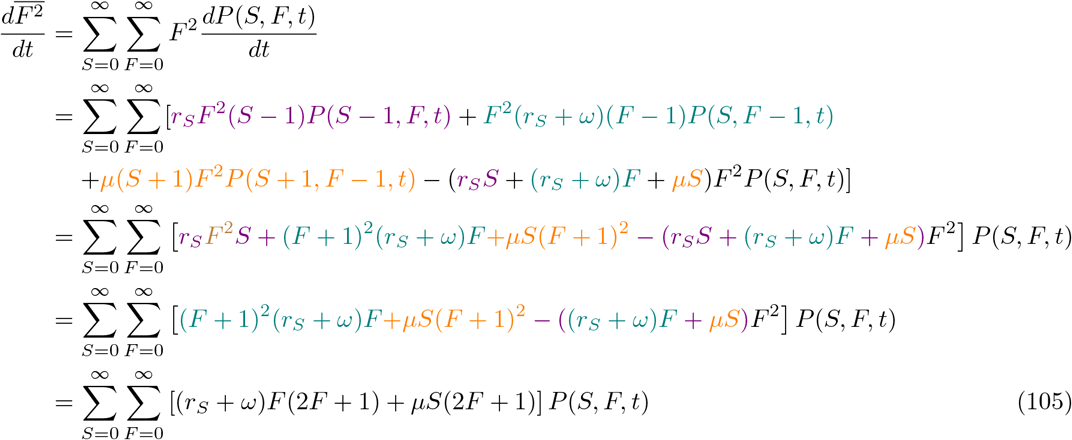

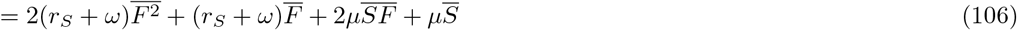

which is Eq. (17).

#### Equation (18) and (20)

Using integration factor 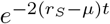 and Eq. (16), we have:

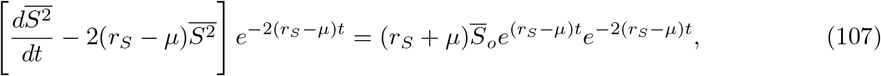

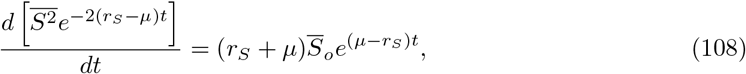

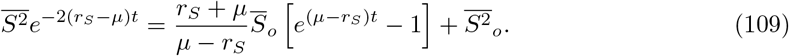

Since *µ ≪ r*_*S*_, we have

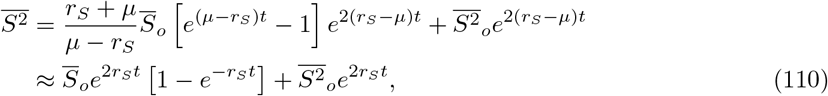

where 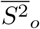 is the expected *S*^2^ at time 0. For Eq. (20),

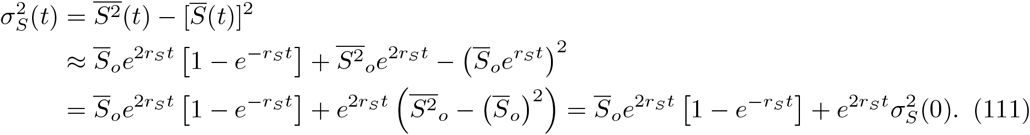

#### Equation (22) and (23)

Since 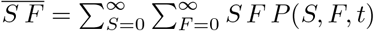,

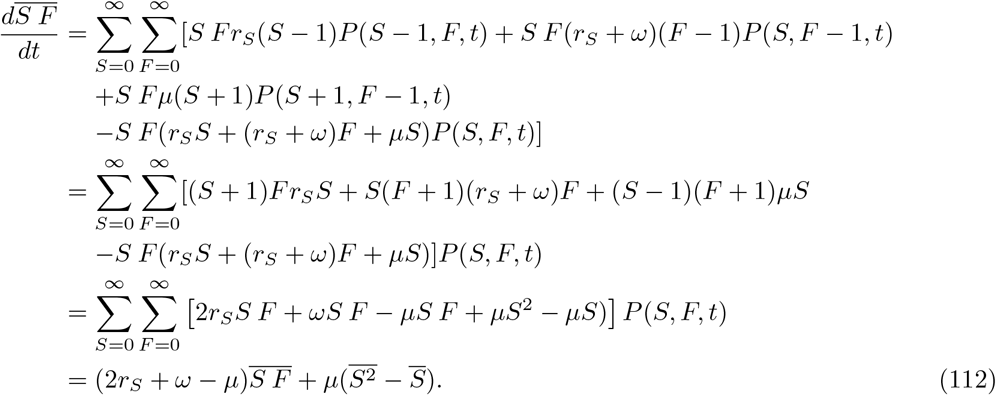

We can solve this, again using the integration factor technique above:

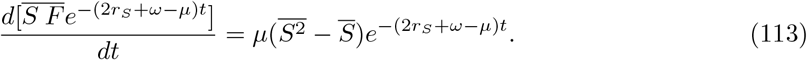

Thus, we have

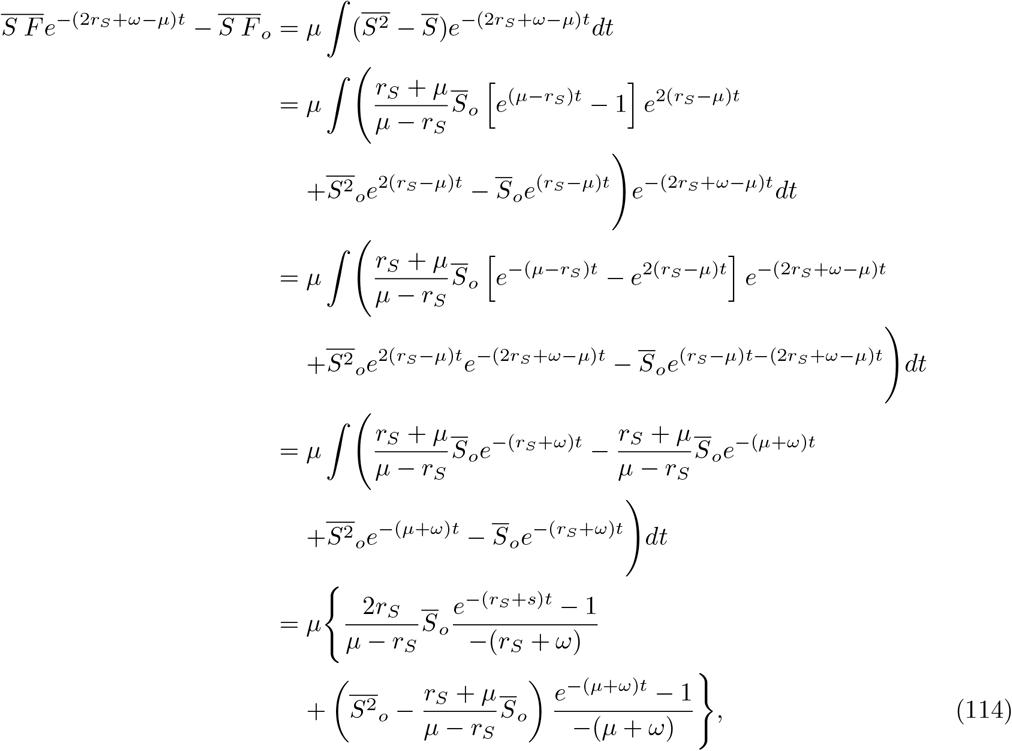

which results in

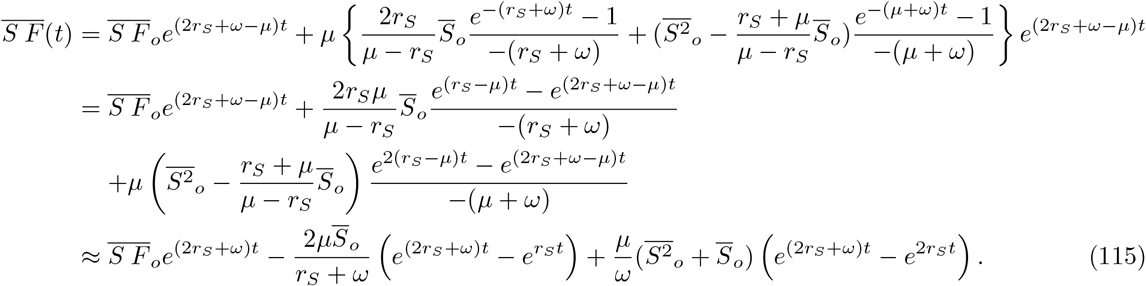

#### Equation (24)

From Eq. (17), we have

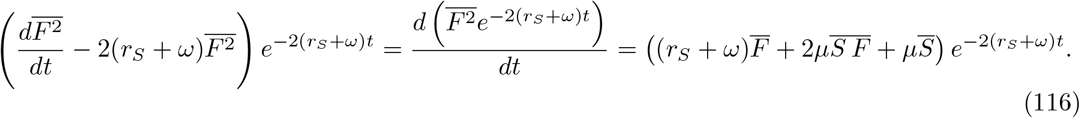

The right-hand side becomes

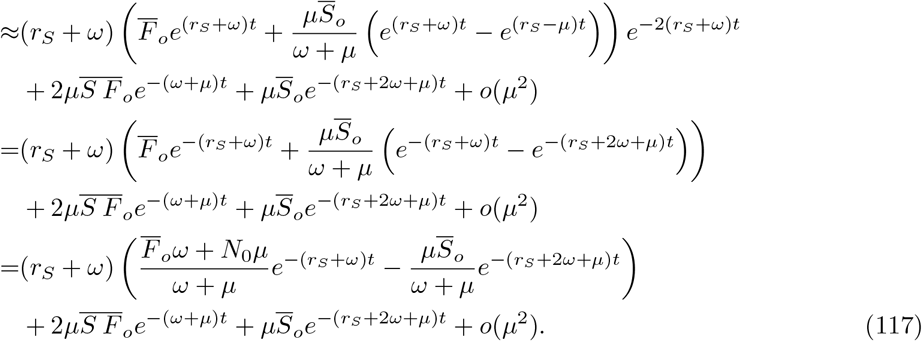

Note that we have checked that the second and third terms of 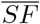 can be ignored after we compare the full calculation with this simpler version. Integrate both sides:

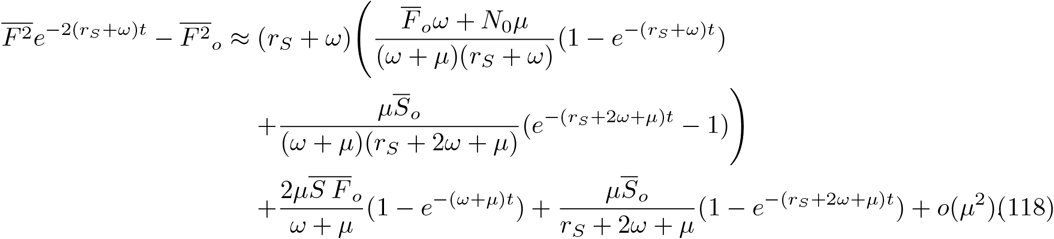

Then, we have

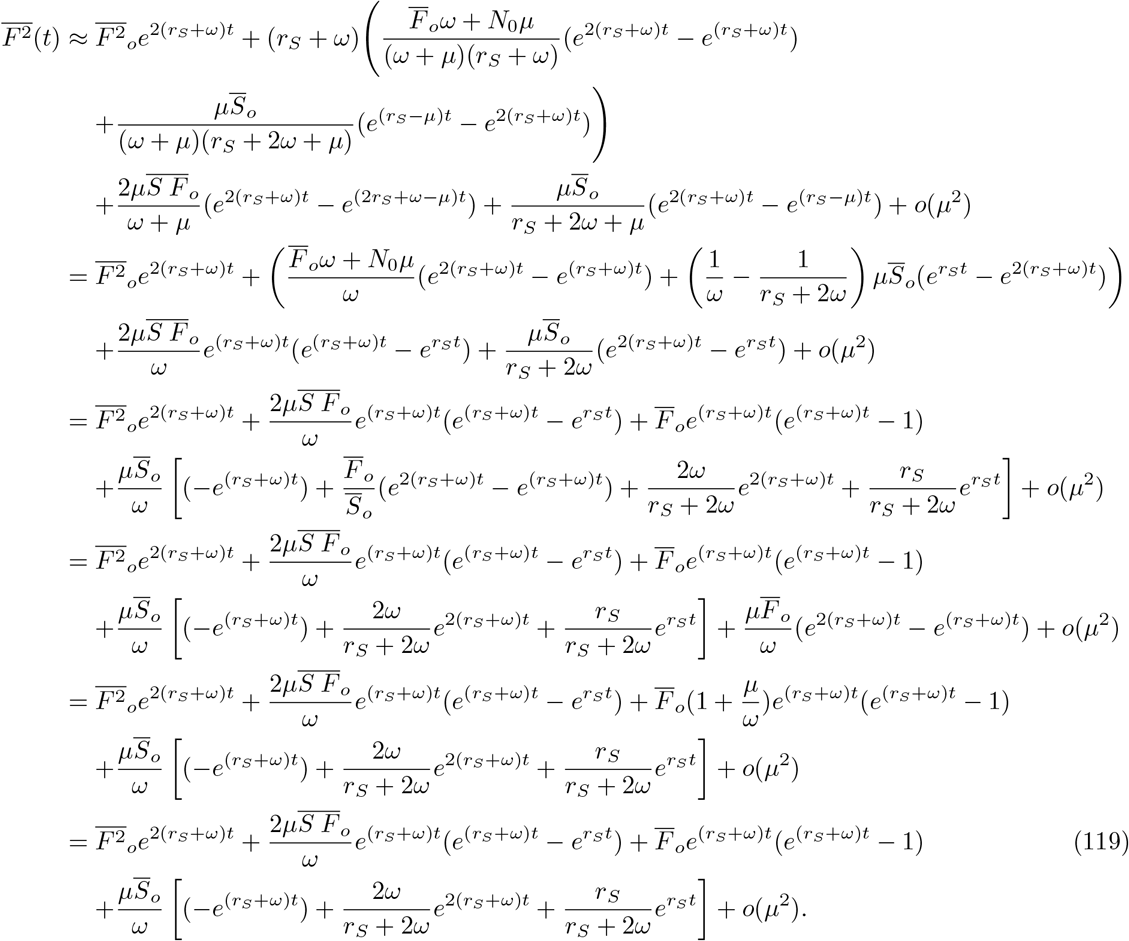

#### Equation (25)

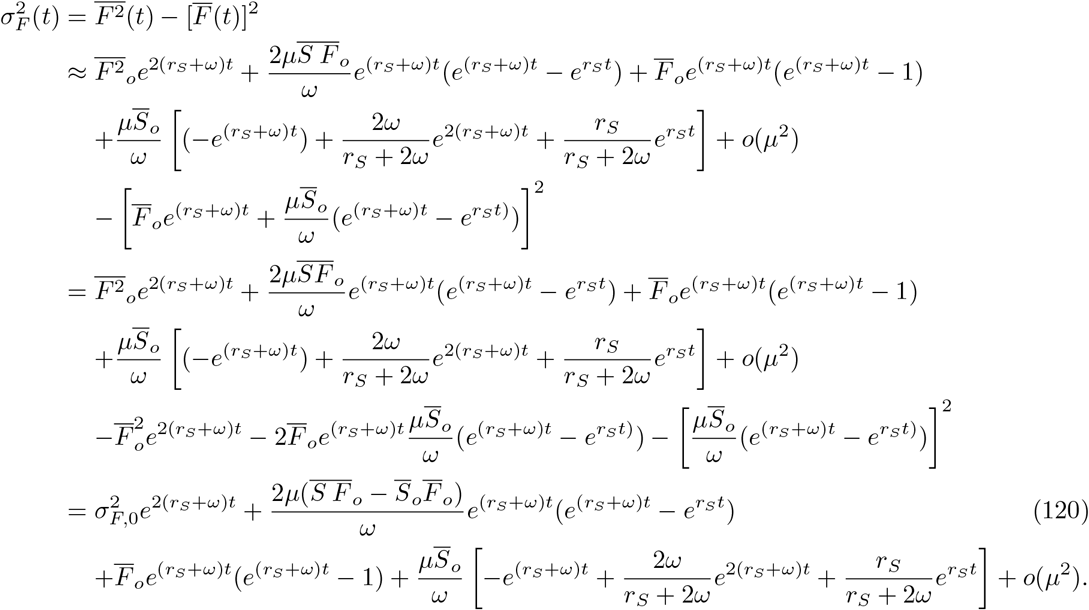

#### Equations (32)-(34)

To derive this equation, we use the fact that 1*/*(1 + *x*) ∼1 − *x* for small *x*. We will omit (*t*) for simplicity. Also note that we are considering relatively large populations so that the standard deviation is much smaller than the mean.

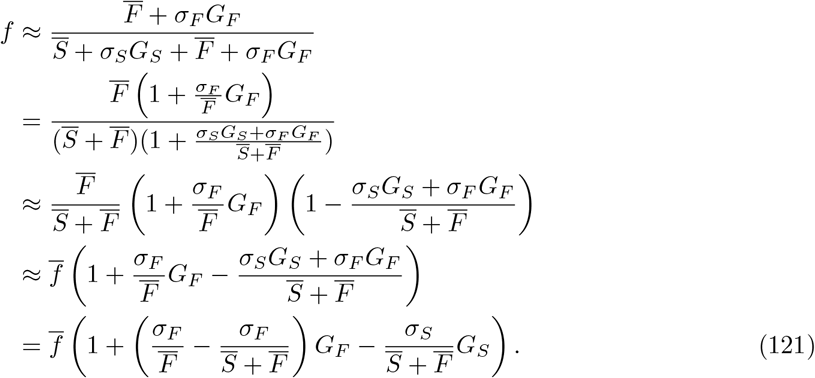

Recall that if *A* ∼ 𝒩 (*µ*_*A*_, *σ*_*A*_), *B* ∼ 𝒩 (*µ*_*B*_, *σ*_*B*_), then 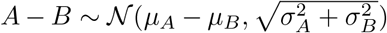.Thus, *f* is distributed as a Gaussian with the mean of

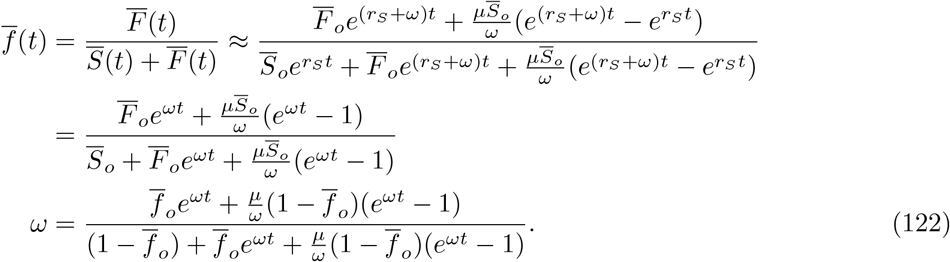

Note that the initial value of mean 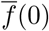 is equal to the mean of the binomial distribution Eq. (29), 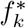.The variance is

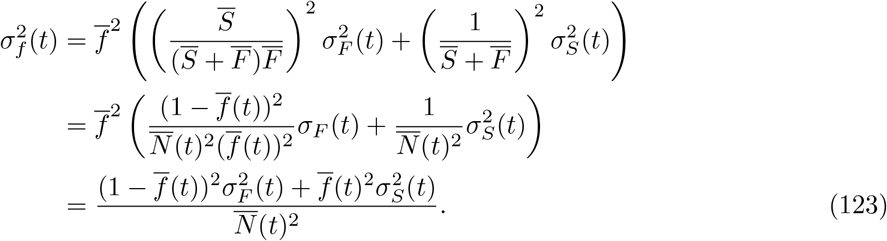

## References

[1] Woo, S., Song, I. & Cha, H. J. Fast and facile biodegradation of polystyrene by the gut microbial flora of Plesiophthalmus davidis larvae. Applied and Environmental Microbiology 86, e01361–20 (2020).

[2] Sun, J., Prabhu, A., Aroney, S. T. & Rinke, C. Corrigendum: Insights into plastic biodegradation: community composition and functional capabilities of the superworm (Zophobas morio) microbiome in styrofoam feeding trials. Microbial Genomics 8 (2022).

[3] Bober, J. R., Beisel, C. L. & Nair, N. U. Synthetic biology approaches to engineer probiotics and members of the human microbiota for biomedical applications. Annual Review of Biomedical Engineering 20, 277–300 (2018).

[4] Wang, E.-X., Ding, M.-Z., Ma, Q.Dong, X.-T. & Yuan, Y.-J. Reorganization of a synthetic microbial consortium for one-step vitamin c fermentation. Microbial Cell Factories 15, 21 (2016).

[5] Goodnight, C. J. Experimental studies of community evolution i: The response to selection at the community level. Evolution 44, 1614–1624 (1990).

[6] Goodnight, C. J. Experimental studies of community evolution ii: The ecological basis of the response to community selection. Evolution 44, 1625–1636 (1990).

[7] Swenson, W., Wilson, D. S. & Elias, R. Artificial ecosystem selection. Proceedings of the National Academy of Sciences 97, 9110–9114 (2000).

[8] Swenson, W., Arendt, J. & Wilson, D. S. Artificial selection of microbial ecosystems for 3-chloroaniline biodegradation. Environmental Microbiology 2, 564–571 (2000).

[9] Blouin, M., Karimi, B., Mathieu, J. & Lerch, T. Z. Levels and limits in artificial selection of communities. Ecology Letters 18, 1040–1048 (2015).

[10] Panke-Buisse, K., Poole, A. C., Goodrich, J. K., Ley, R. E. & Kao-Kniffin, J. Selection on soil microbiomes reveals reproducible impacts on plant function. The ISME Journal 9, 980–989 (2015).

[11] Panke-Buisse, K., Lee, S. & Kao-Kniffin, J. Cultivated sub-populations of soil microbiomes retain early flowering plant trait. Microbial Ecology 73, 394–403 (2017).

[12] Jochum, M. D., McWilliams, K. L., Pierson, E. A. & Jo, Y.-K. Host-mediated microbiome engineering (hmme) of drought tolerance in the wheat rhizosphere. PLoS One 14, e0225933 (2019).

[13] Wright, R. J., Gibson, M. I. & Christie-Oleza, J. A. Understanding microbial community dynamics to improve optimal microbiome selection. Microbiome 7, 1–14 (2019).

[14] Raynaud, T., Devers, M., Spor, A. & Blouin, M. Effect of the reproduction method in an artificial selection experiment at the community level. Frontiers in Ecology and Evolution 7, 416 (2019).

[15] Jigyasa Arora, A. S. M., Margaret A. Mars Brisbin. Effects of microbial evolution dominate those of experimental host-mediated indirect selection. PeerJ 8, e9350 (2020).

[16] Chang, C.-Y., Osborne, M. L., Bajic, D. & Sanchez, A. Artificially selecting bacterial communities using propagule strategies. Evolution 74, 2392–2403 (2020).

[17] Mueller, U. G. et al. Artificial selection on microbiomes to breed microbiomes that confer salt tolerance to plants. mSystems 6, e01125–21 (2021).

[18] Jacquiod, S. et al. Artificial selection of stable rhizosphere microbiota leads to heritable plant phenotype changes. Ecology Letters 25, 189–201 (2022).

[19] Raynaud, T., Devers-Lamrani, M., Spor, A. & Blouin, M. Community diversity determines the evolution of synthetic bacterial communities under artificial selection. Evolution 76, 1883–1895 (2022).

[20] Penn, A. “Modelling artificial ecosystem selection: A preliminary investigation”. In Banzhaf, W., Ziegler, J., Christaller, T., Dittrich, P. & Kim, J.T. (eds.) Advances in Artificial Life, 659–666 (Springer Berlin Heidelberg, Berlin, Heidelberg, 2003).

[21] Penn, A. & Harvey, I. “The role of non-genetic change in the heritability, variation, and response to selection of artificially selected ecosystems”. In Pollack, J., Bedau, M. A., Husbands, P., Watson, R.A. & Ikegami, T. (eds.) Artificial Life IX: Proceedings of the Ninth International Conference on the Simulation and Synthesis of Artificial Life, vol. 9, 352 (MIT Press, 2004).

[22] Williams, H. T. P. & Lenton, T. M. Artificial selection of simulated microbial ecosystems. Proceedings of the National Academy of Sciences 104, 8918 (2007).

[23] Xie, L., Yuan, A. E. & Shou, W. Simulations reveal challenges to artificial community selection and possible strategies for success. PLoS Biology 17, e3000295 (2019).

[24] Doulcier, G., Lambert, A., De Monte, S. & Rainey, P. B. Eco-evolutionary dynamics of nested darwinian populations and the emergence of community-level heredity. eLife 9, e53433 (2020).

[25] Xie, L. & Shou, W. Steering ecological-evolutionary dynamics to improve artificial selection of microbial communities. Nature Communications 12, 6799 (2021).

[26] Chang, C.-Y. et al. Engineering complex communities by directed evolution. Nature Ecology & Evolution 5, 1011 (2021).

[27] Fraboul, J., Biroli, G. & De Monte, S. Artificial selection of communities drives the emergence of structured interactions. Journal of Theoretical Biology 571, 111557 (2023).

[28] Lalejini, A., Dolson, E., Vostinar, A. E. & Zaman, L. Artificial selection methods from evolutionary computing show promise for directed evolution of microbes. eLife 11, e79665 (2022).

[29] Zaccaria, M., Sandlin, N., Soen, Y. & Momeni, B. Partner-assisted artificial selection of a secondary function for efficient bioremediation. iScience 26, 107632 (2023).

[30] Vessman, B., Guridi-Fernández, P., Arias-Sánchez, F. I. & Mitri, S. Novel artificial selection method improves function of simulated microbial communities. bioRxiv [preprint] (2023). 10.1101/2023.01.08.523165 (accessed 18 December 2023).

[31] Arias-Sánchez, F. I., Vessman, B., Haym, A., Alberti, G. & Mitri, S. Artificial selection improves pollutant degradation by bacterial communities. Nature Communications 15, 7836 (2024).

[32] Xie, L., Yuan, A. E. & Shou, W. A quantitative genetics framework for understanding the selection response of microbial communities. bioRxiv [preprint] (2023). 10.1101/2023.10.24.563725 (accessed 28 December 2023).

[33] Thomas, J. L., Rowland-Chandler, J. & Shou, W. Artificial selection of microbial communities: what have we learnt and how can we improve? Current Opinion in Microbiology 77, 102400 (2024).

[34] Rainey, P. B. Major evolutionary transitions in individuality between humans and ai. Philosophical Transactions of the Royal Society B 378, 20210408 (2023).

[35] Philip, J. The function inverfc theta. Australian Journal of Physics 13, 13 (1960).

[36] Ikeda, Y., Grabowski, B. & Körmann, F. mpltern 0.3.0: ternary plots as projections of Matplotlib (2019). URL 10.5281/zenodo.3528355.

[37] Gillespie, D. T. Approximate accelerated stochastic simulation of chemically reacting systems. The Journal of Chemical Physics 115, 1716–1733 (2001).

[38] Cao, Y., Gillespie, D. T. & Petzold, L. R. Efficient step size selection for the tau-leaping simulation method. The Journal of Chemical Physics 124, 044109 (2006).

[39] Gumbel, E. J. Statistics of extremes (Columbia university press, 1958).

## References

[1] Gillespie, D. T. Approximate accelerated stochastic simulation of chemically reacting systems. The Journal of Chemical Physics 115, 1716–1733 (2001).

[2] Cao, Y., Gillespie, D. T. & Petzold, L. R. Efficient step size selection for the tau-leaping simulation method. The Journal of Chemical Physics 124, 044109 (2006).

[3] Zheng, Q. Progress of a half century in the study of the luria–delbrück distribution. Mathematical Biosciences 162, 1–32 (1999).

[4] Gumbel, E. J. Statistics of extremes (Columbia university press, 1958).

[5] Philip, J. The function inverfc theta. Australian Journal of Physics 13, 13 (1960).

[6] Xie, L., Yuan, A. E. & Shou, W. Simulations reveal challenges to artificial community selection and possible strategies for success. PLoS Biology 17, e3000295 (2019).

[7] Ikeda, Y., Grabowski, B. & Körmann, F. mpltern 0.3.0: ternary plots as projections of Matplotlib (2019). Available at 10.5281/zenodo.3528355.

